# A tailored *in vivo* CRISPR screen identifies *BAP1* as a potent tumor suppressor of soft tissue sarcoma

**DOI:** 10.64898/2025.12.01.691734

**Authors:** Jianguo Huang, Xingliang Liu, Warren Floyd, William Haugh, Zhaoyu Sun, Melissa J. Kasiewicz, Yaping Wu, Brian Piening, John T. Welle, Wesley K. Rosales, Venkatesh Rajamanickam, So Young Kim, Eric Xu, Lixia Luo, Yan Ma, Rutulkumar Patel, Ziqiang Zhang, Brady Bernard, William L. Redmond, Walter J. Urba, R Bryan Bell, David G. Kirsch

**Affiliations:** Earle A. Chiles Research Institute, Providence Cancer Institute, Portland, OR 97213; Department of Radiation Oncology, Duke University Medical Center, Durham, NC 27710; Department of Molecular Genetics and Microbiology, Duke University, Durham, NC 27710; Department of Respiratory and Critical Care Medicine, Shanghai Pudong Hospital, Fudan University Pudong Medical Center, Pudong Hospital of Fudan University, Shanghai, China; Department of Pharmacology and Cancer Biology, Duke University Medical Center, Durham, NC 27710; Radiation Medicine Program, Princess Margaret Cancer Center, University Health Network, Toronto, ON, Canada; Department of Radiation Oncology, University of Toronto, Toronto, ON, Canada; Department of Medical Biophysics, University of Toronto, Toronto, ON, Canada

## Abstract

Undifferentiated pleomorphic sarcoma (UPS) is one of the most common soft tissue sarcomas (STS) in adults. Despite decades of research, therapeutic advancements for STS, including UPS, have remained limited. The genetic complexity of UPS, characterized by the absence of recurrent driver oncogene mutations, has hindered the development of effective targeted therapies beyond conventional chemotherapy and immunotherapy. To address this challenge, we conducted a customized *in vivo* CRISPR/Cas9 screen in mice to systematically identify potential tumor suppressors involved in UPS development. Our screen revealed BRCA1-associated protein 1 (Bap1) as a potent tumor suppressor in STS. Using total RNA sequencing, multiplex immunohistochemistry, and flow cytometry, we found that *Bap1*-deficient mouse sarcomas exhibit significant immune suppression. Further analysis indicated that polo-like kinase 1 (Plk1) is essential for the survival of *Bap1*-deficient sarcomas. Treatment with volasertib, a Plk1 inhibitor, markedly inhibited tumor growth in both syngeneic and spontaneous mouse models of *Bap1*-loss sarcoma. In conclusion, our findings suggest that PLK1 inhibition, or combined with immunotherapy, may represent a promising targeted therapeutic strategy for tumors lacking *BAP1*.

## Introduction

Soft tissue sarcomas (STS) are rare mesenchymal malignancies that originate from diverse connective tissues ^1^. STS are further subclassified into over 75 subtypes, posing significant challenge of developing novel therapies for each STS subtype. Undifferentiated pleomorphic sarcoma (UPS) is one of the most common sarcoma subtypes diagnosed in adults. As a high-grade STS, approximately 50% of patients with UPS develop lung metastasis with a median survival of less than 15 months ^2,3^. For decades, little therapeutic progress has been made in the treatment of STS. Anthracycline (doxorubicin)-based chemotherapy has remained the standard chemotherapy treatment of high-grade STS, including metastatic UPS, for over 40 years ^4^. However, unlike other cancers driven by well-characterized genetic mutations, UPS lacks recurrent oncogenic drivers, limiting the development of targeted therapies ^5,6^. Identifying genetic drivers in UPS could accelerate the discovery of effective therapeutic strategies for this challenging disease.

Research on human UPS is constrained by the limited availability of biospecimens and reagents, which hampers basic investigations and therapeutic testing. The genetic complexity and absence of well-defined driver mutations further complicate the development of targeted treatments. Genetically engineered mouse models that recapitulate subsets of human UPS are essential tools for studying disease biology and evaluating novel therapies. Several primary mouse models have been established, including primary tumors driven by activation of conditional oncogenic *Kras*^G12D^ and deletion of *Trp53* (KP) or *Cdkn2a* (KI), deletion of both *Trp53* and *Pten* (PP), deletion of *NF1* with *Trp53* (NP) or *Cdkn2a* (NI), and deletion of both *Trp53* and *Rb1* (PR) ^7–11^. These models have been instrumental in elucidating mechanisms of sarcomagenesis, metastasis, and immune evasion, and in guiding immunotherapeutic strategies ^3,10,12–16^. However, the relevance of these models to human UPS is limited. For instance, *KRAS*^G12D^ as an oncogenic driver that are rarely found in human UPS, while *NF1* and *PTEN* are tumor suppressor drivers mutated only in a small subset of human UPS. The molecular pathogenesis of most human UPS remains unknown. After *TP53*, *RB1* is the second most frequently mutated gene in UPS, often through loss-of-function mutations or copy number alterations. Yet, compared to KP models, mice with spatial and temporal deletion of *Trp53* and *Rb1* develop sarcomas less frequently and with delayed onset ^10^. Whether additional genetic alterations could cooperate with *Trp53* and *Rb1* loss to accelerate tumorigenesis remains unexplored.

BRCA1-associated protein 1 (BAP1) is a tumor suppressor frequently mutated in uveal melanoma, malignant mesothelioma, cutaneous melanoma, renal cell carcinoma (RCC), and other malignancies, including STS ^17^. Germline mutations in *BAP1* define a tumor predisposition syndrome (BAP1-TPDS), and STS has been reported in affected individuals ^18^. Transgenic mouse models have demonstrated that *Bap1* deletion can increase malignancy of mesothelioma, RCC, and other cancers ^19–22^. Additionally, it is suggested that *BAP1* mutations are a late stage event during cancer progression ^23^. Furthermore, the role of *BAP1* in STS remains uncharacterized. *BAP1* encodes a deubiquitylase that forms the polycomb repressive deubiquitylase (PR-DUB) complex, which reverses monoubiquitylation of histone H2A at lysine 119 (H2AK119ub1), a modification catalyzed by polycomb repressive complex 1 (PRC1) ^24^. Loss of *Bap1* leads to diffuse accumulation of H2AK119ub1, which results in global chromatin compaction. This compaction activates the DNA damage response and impairs DNA repair, ultimately compromising cell survival ^25^. Thus, *BAP1* is thought to suppress tumorigenesis through epigenetic regulation during cancer progression ^26–28^.

Polo-like kinase 1 (PLK1) is a serine/threonine kinase essential for mitotic progression, regulating centrosome maturation, spindle assembly, and chromosome segregation ^29–31^. *PLK1* dysregulation has been implicated in various cancers, and its overexpression drives tumorigenesis in transgenic mouse models, including STS ^32,33^. These findings support *PLK1* as an oncogenic driver in sarcomagenesis. However, analysis of STS samples in The Cancer Genome Atlas (TCGA) reveals no recurrent copy number alterations or mutations in *PLK1*. In this study, we investigate the mechanisms underlying *PLK1* induction and evaluate its potential as a therapeutic target in *BAP1*-deficient sarcomas.

## Results

### Intramuscular injection of a CRISPR sgRNA library induces spontaneous sarcomas in mice

To investigate the tumor suppressor function of genes frequently altered in human undifferentiated pleomorphic sarcoma (UPS), we employed a direct in vivo autochthonous screening strategy. This approach enables targeted mutagenesis within the native skeletal muscle microenvironment of mice ^34^. Using data from The Cancer Genome Atlas (TCGA) and the Genomics Evidence Neoplasia Information Exchange (GENIE), we identified the top 35 genes with loss-of-function mutations or copy number alterations (CNAs) in human UPS and myxofibrosarcoma (MFS), which share similar genetic profiles (**Supplementary Table 1**), along with tumor protein p53 (*TP53*) and cyclin-dependent kinase inhibitor 2A (*CDKN2A*) ^35^. We designed a custom single-guide RNA (sgRNA) library targeting these 35 genes (excluding *Trp53* and *Cdkn2a*), with four sgRNAs per gene sourced from the mouse Brie library (**Supplementary Table 2**) ^36^. Additionally, we included sgRNAs targeting 5 housekeeping genes: Ribosomal protein L7 (*Rpl7*), Ribosomal proteins 19 (*Rps19*), Ribosomal protein L22 (*Rpl22*), Ribosomal proteins 18 (*Rps18*), and Ribosomal proteins 11 (*Rps11*) ^34^. The customized sgRNA library also included 8 nontargeting control sgRNAs from the mouse Brie library (**Supplementary Table 2**). Given that *TP53* is mutated in approximately 65% of human UPS and MFS (**Supplementary Figure 1**), and that deletion of *Cdkn2a* can substitute for *Trp53* loss to induce sarcomagenesis in conjunction with *Kras*^G12D^ ^37^, we cloned this customized sgRNA library into 3 different plasmid backbones: 1) negative control sgRNA expressing plasmid (pX334-Negative sgRNA-library); 2) *Trp53* sgRNA expressing plasmid (pX334-Trp53 sgRNA-library); and 3) *Cdkn2a* sgRNA expressing plasmid (pX334-Cdkn2a sgRNA-library) (**Figure 1A**). The *Trp53* sgRNA and *Cdkn2a* sgRNA in these constructs have been confirmed to induce robust and specific editing in the targeting loci, according to previous studies ^37,38^. All three plasmid libraries were sequenced to confirm full coverages of all sgRNAs (**Supplementary Figure 2**). We utilized an in vivo electroporation (IVE) method to deliver these plasmids into the gastrocnemius muscle of Cas9-expressing mice, enabling CRISPR/Cas9-mediated mutagenesis of tumor suppressor genes and generation of UPS-like tumors ^11,37^. Only the pX334-Trp53 sgRNA-library induced tumors in both Rosa26 ^LoxP-Cas9/LoxP-Cas9^ and Rosa26 ^LoxP-Cas9/+^ mice (**Figure 1B**, **Supplementary Figure 3**). In an independent cohort, we confirmed that the pX334-Trp53 sgRNA-library induced tumors in Rosa26 ^LoxP-Cas9/LoxP-Cas9^ mice, whereas the pX334-Trp53 sgRNA alone (lacking sgRNAs for other tumor suppressors) failed to induce tumors (**Figure 1C**). These results suggest that mutation of at least one additional gene from the library is required to cooperate with *Trp53* loss in driving tumorigenesis.

**Figure 1.**
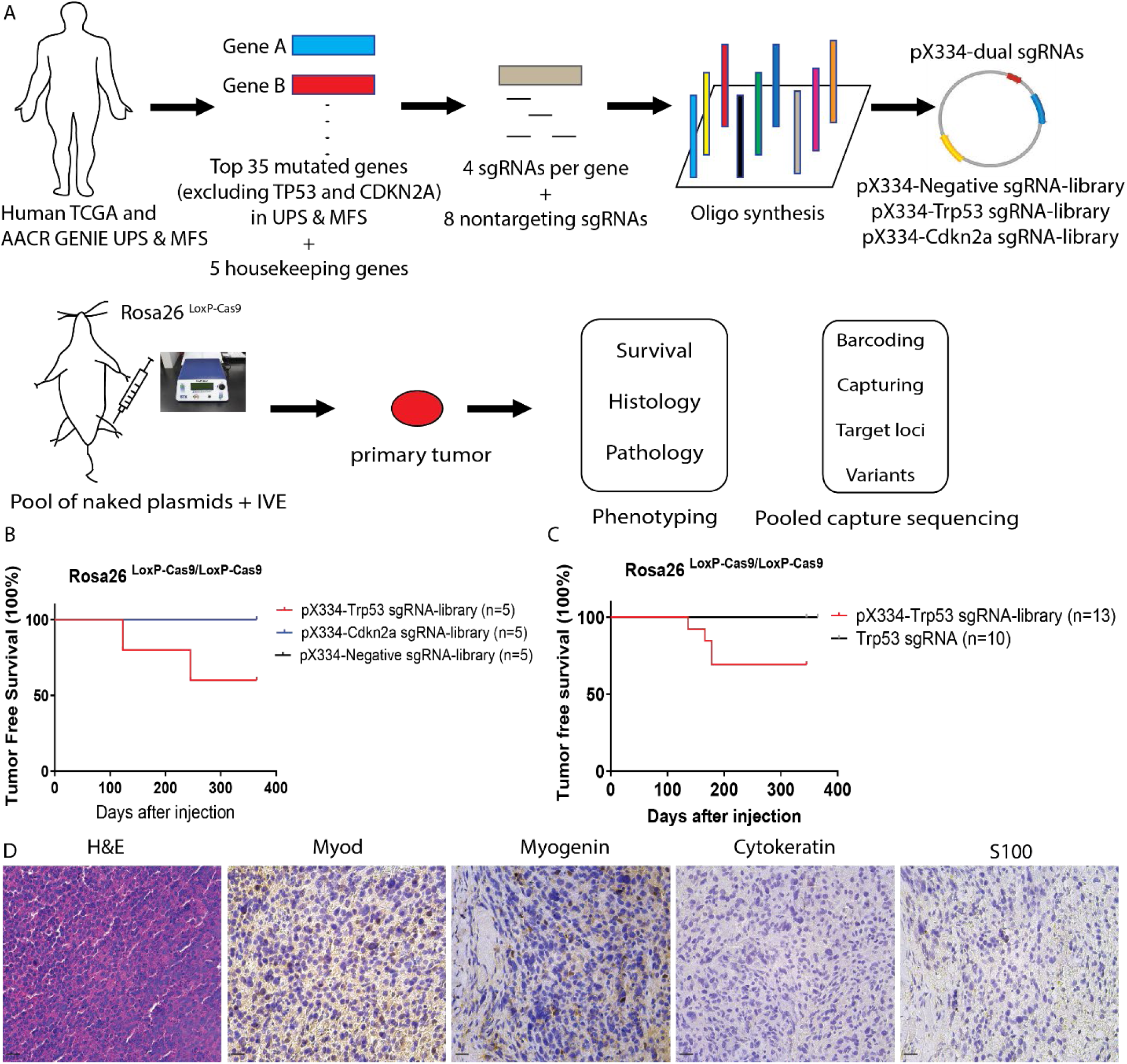
*In vivo* CRISPR/Cas9 screen identifies tumor suppressors of UPS in Rosa26 ^LoxP-Cas9^ mice. (A) Schematic of the direct *in vivo* CRISPR/Cas9 screening strategy. (B) Two primary tumors developed in Rosa26 LoxP-Cas9/LoxP-Cas9 mice following intramuscular delivery of the pX334-Trp53 sgRNA-library (n = 5), while no tumors formed in mice receiving the pX334-Negative sgRNA-library or pX334-Cdkn2a sgRNA-library (n = 5 each). (C) In a separate cohort, five tumors developed in mice injected with the pX334-Trp53 sgRNA-library (n = 13), but no tumors formed in mice injected with pX334-Trp53 sgRNA alone (n = 10). (D) Representative H&E and IHC staining (MyoD, Myogenin, Cytokeratin, S100) of a tumor induced by the pX334-Trp53 sgRNA-library. Scale bar = 100 µm.

To characterize the tumors induced by the pX334-Trp53 sgRNA-library, we performed hematoxylin and eosin (H&E) staining and immunohistochemistry (IHC) on sections from 10 tumors using antibodies against Cytokeratin, MyoD, Myogenin, and S100 (**Figure 1D**). A blinded pathological review determined that all tumors were poorly differentiated, negative for Cytokeratin and S100, and weak or negative for MyoD and Myogenin—consistent with UPS or myogenic UPS phenotypes.

### Targeted capture sequencing identifies multiple tumor suppressor candidates from the Trp53-library-induced tumors

We generated a total of 14 tumors in mice using the pX334-Trp53 sgRNA-library alone, and an additional three tumors following co-delivery of the pX334-Trp53 sgRNA-library with a *Pten* sgRNA (**Supplementary Figure 4**). To validate CRISPR/Cas9-mediated gene editing using our in vivo electroporation (IVE) method, we injected the **pX334-Trp53 sgRNA-library** into the gastrocnemius muscle of two mice. Four weeks post-injection, the muscle tissues were harvested, and genomic DNA was extracted from both tumors (S1–S17) and injected muscle tissues (S18–S19) for targeted capture sequencing ^34^.

Bioinformatic analysis revealed CRISPR/Cas9-mediated editing in *Cysltr2* (S18) and *Pask* (S19) in the injected muscle tissues. However, editing was not detected in most targeted genes, including *Trp53*, in these samples. This may be due to the absence of transformed cells harboring *Trp53* mutations in the harvested tissues, or the inability of our sequencing method to detect low-frequency edits in underrepresented mutant cells. Interestingly, *CYSLTR2* ranks among the top three genes with loss-of-function mutations or CNAs in human UPS and MFS, following *TP53* and *RB1*. *PASK* also shows the highest mutation frequency in UPS and MFS compared to other cancers in the TCGA database. These findings confirm that our IVE method can induce gene editing in vivo using pooled plasmid libraries (**Figure 2A**).

**Figure 2.**
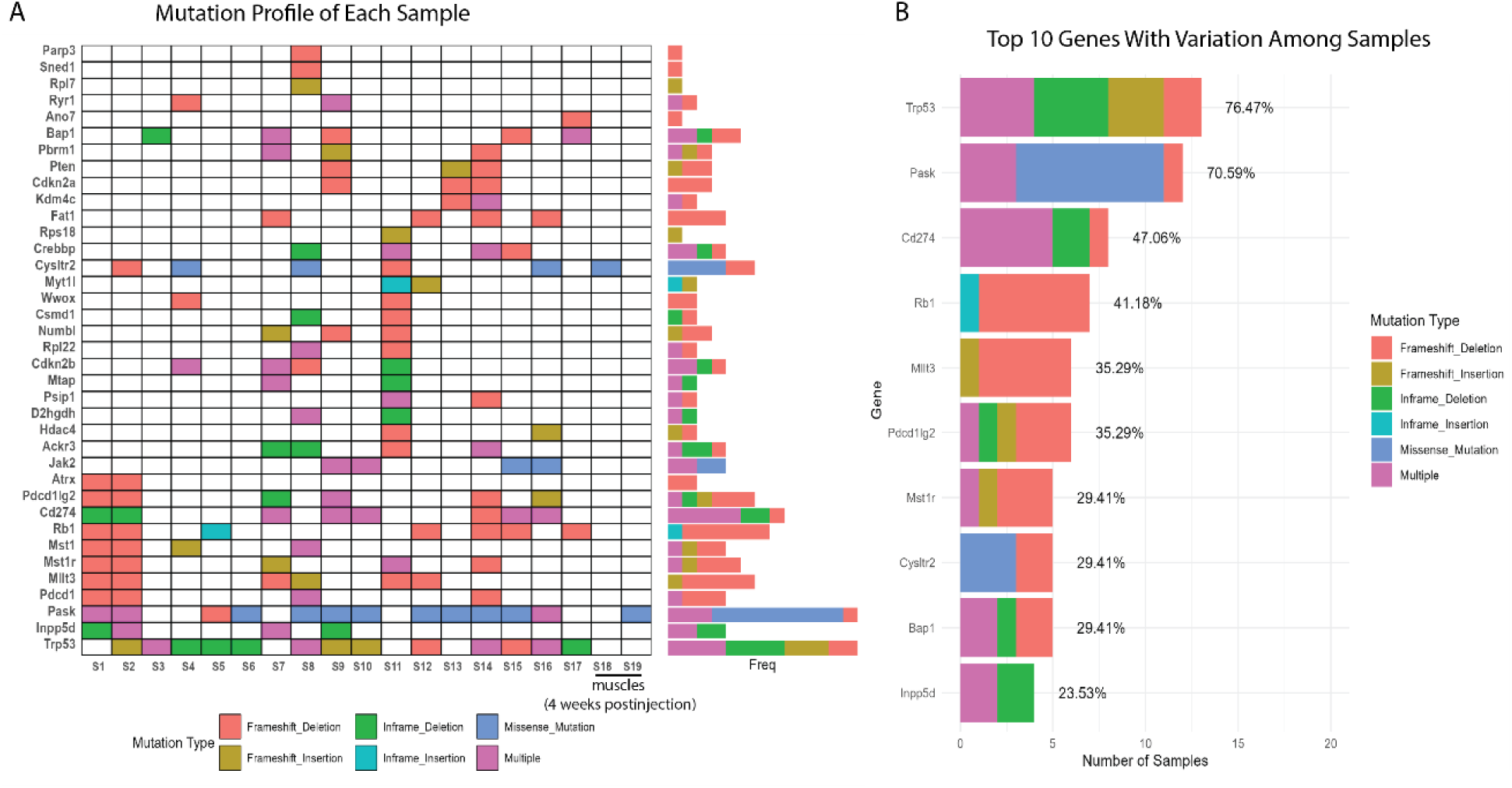
Targeted capture sequencing of tumors induced by the pX334-Trp53 sgRNA-library. (A) Mutation profiles from targeted capture sequencing of injected muscle tissues (S18–S19) and tumors (S1–S17), including three tumors with additional *Pten* mutations (S9, S13, S14). (B) Top 10 most frequently mutated genes across all tumors.

In tumors induced by co-delivery of the pX334-Trp53 sgRNA-library and *Pten* sgRNA, editing of *Pten* was detected exclusively in those samples (**Figure 2A**, **Supplementary Figure 4**). Editing of *Trp53* was detected in nearly all tumors, although four tumors (S1, S7, S11, S13) showed no detectable edits by targeted capture sequencing. To further investigate, we performed T7 endonuclease I (T7E1) assays and Sanger sequencing on genomic DNA from these tumors. In tumor S1, we confirmed 82% insertions and deletions (Indels) at the *Trp53* locus, including a three-nucleotide deletion in one allele (**Supplementary Figure 5A and B**). It is possible that the other allele harbors a large deletion undetectable by our method, consistent with previous reports of large CRISPR-induced deletions in *Trp53* ^11^. Due to limitations in Sanger sequencing sensitivity ^39^, we performed next-generation sequencing (NGS) on tumors S7, S11, and S13. We detected approximately 9% Indels in S7, 11% in S11, and 12% in S13, with multiple Indel variants per sample (**Supplementary Table 3 and Supplementary Figure 5C**). The relatively low Indel percentages may reflect the presence of normal cells mixed with tumor tissue.

Beyond *Trp53*, other frequently mutated genes in the tumors included *Mllt3* (also known as *Af9*), *Fat1*, *Bap1*, and *Rb1* (**Figure 2A and 2B**). Notably, only *Trp53* and *Bap1* were mutated in tumor S3. According to TCGA data, *BAP1* mutations or CNAs are predominantly found in UPS among all STS subtypes (**Supplementary Figure 1**). Tumor S6 harbored mutations in *Trp53* and *Pask*. To further investigate the role of these candidate tumor suppressors, we individually evaluated their ability to cooperate with *Trp53* mutation in driving sarcoma development.

### Validation of tumor suppressor genes cooperating with *Trp53* mutation to drive sarcoma development in mice

To validate candidate tumor suppressor genes identified from the CRISPR screen, we performed intramuscular injections of plasmids expressing sgRNAs targeting individual genes along with *Trp53* sgRNA into Rosa26 ^LoxP-Cas9/LoxP-Cas9^ mice. Tumor formation was observed in mice electroporated with plasmids targeting *Trp53* and *Bap1*, as well as those targeting *Trp53* and *Rb1* (**Figure 3A**). Tumors also formed in mice injected with plasmids targeting *Trp53* and *Pask*, although with low penetrance (2 out of 10 mice; data not shown). This low penetrance is currently under further investigation and may require validation in a larger cohort and in a transgenic model of Trp53 ^FL/FL^ and Pask ^FL/FL^. In contrast, no tumors were observed in mice injected with plasmids targeting *Trp53* in combination with *Cysltr2*, *Mst1r*, *Mllt3*, *Crebbp*, *Pbrm1*, or *Fat1* (**Supplementary Figure 6**). We next compared tumor growth rates between different genotypes. Tumors generated by deletion of *Trp53* and *Rb1* (referred to as PR CRISPR tumors) and those generated by deletion of *Trp53* and *Bap1* (PB CRISPR tumors) showed no significant difference in time to quadrupling of tumor volume (**Figure 3B and 3C**).

**Figure 3.**
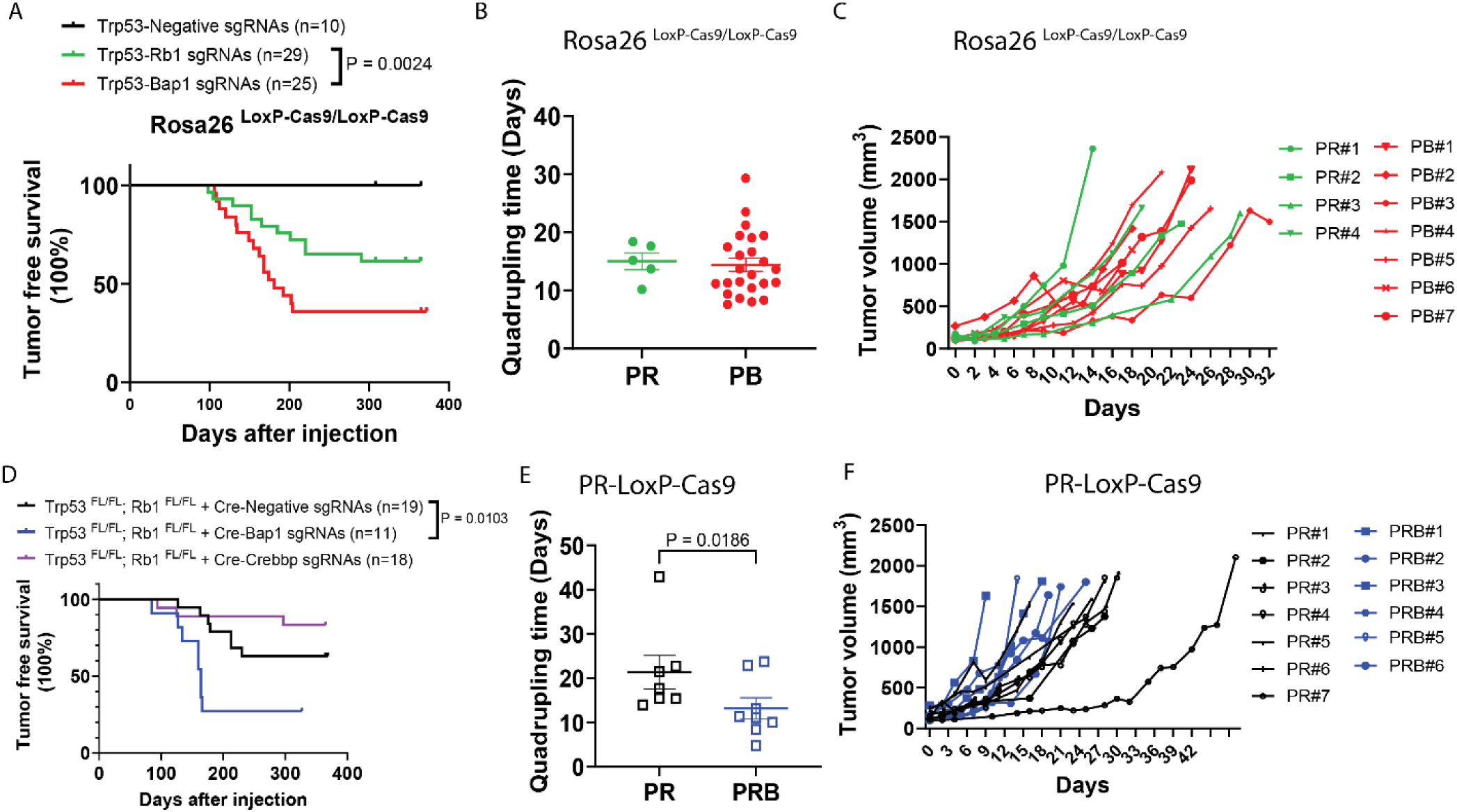
Validation of *Bap1* as a potent tumor suppressor in UPS. (A) IVE of plasmids expressing sgRNAs targeting *Trp53* and *Bap1* or *Trp53* and *Rb1* induced tumors in Rosa26 ^LoxP-Cas9/LoxP-Cas9^ mice. (B) Tumor quadrupling times were similar between PR CRISPR and PB CRISPR tumors. (C) Growth curves of representative PR and PB CRISPR tumors. (D) IVE of Cre recombinase and sgRNA targeting *Bap1* (n = 11) induced significantly higher tumor penetrance compared to Cre alone or Cre + sgRNA targeting *Crebbp* (3 tumors in 18 mice). (E) PRB Cre/CRISPR tumors grew significantly faster than PR Cre tumors. (F) Growth curves of representative PR Cre and PRB Cre/CRISPR tumors.

To further explore genetic cooperation, we investigated whether additional mutations could accelerate tumorigenesis in the context of *Trp53* and *Rb1* loss. Previous studies in small cell lung cancer models have shown that *Crebbp* deletion accelerates tumor formation when combined with *Trp53* and *Rb1* loss ^40^. TCGA data also indicate that *BAP1* copy number loss frequently co-occurs with *RB1* loss in human UPS (**Supplementary Figure 1**). Therefore, we generated a transgenic mouse model of Trp53 ^FL/FL^; Rb1 ^FL/FL^; Rosa26 ^LoxP-Cas9/LoxP-Cas9^ (PR-loxP-Cas9 mice) and performed IVE to deliver plasmids expressing Cre recombinase and sgRNAs targeting either no gene, *Bap1*, or *Crebbp*. Targeting *Crebbp* did not affect tumor development initiated by Cre-mediated deletion of *Trp53* and *Rb1* (**Figure 3B**). In contrast, additional mutation of *Bap1* significantly accelerated tumor onset and increased tumor penetrance in PR-loxP-Cas9 mice (**Figure 3B**). We then compared tumor growth rates between PR Cre tumors (deletion of *Trp53* and *Rb1*) and PRB Cre/CRISPR tumors (deletion of *Trp53*, *Rb1*, and *Bap1*). PRB Cre/CRISPR tumors exhibited significantly faster growth, with shorter times to quadrupling of tumor volume (**Figure 3E and 3F**). These findings suggest that *Bap1* loss enhances sarcoma aggressiveness in the context of *Trp53* and *Rb1* deficiency.

Additionally, we observed lung metastases in mice bearing PR CRISPR tumors (1/11), PR Cre tumors (1/2), PB CRISPR tumors (4/16), and PRB Cre/CRISPR tumors (1/6). Multiple metastatic nodules were detected in the lungs of mice with PB CRISPR tumors (**Supplementary Table 4**). Histological analysis via H&E staining confirmed that all tumors were high-grade undifferentiated sarcomas resembling UPS (**Supplementary Figure 7**).

Taken together, these results demonstrate that *Bap1* functions as a potent tumor suppressor in UPS, both in the context of *Trp53* loss alone and in combination with *Rb1* loss.

### High-throughput RNA sequencing reveals immune suppression in *Bap1*-loss mouse sarcomas

To systematically investigate the signaling pathways contributing to sarcoma development, we performed total RNA sequencing on tumors from three genotypes: PB CRISPR (n = 10), PR Cre (n = 5), and PRB Cre/CRISPR (n = 7), along with normal skeletal muscle controls. Bioinformatic analysis and gene set enrichment analysis (GSEA) revealed that PB CRISPR tumors were significantly enriched for oncogenic pathways, including the E2F targets, G2M checkpoint, and MYC targets hallmarks, compared to normal muscle tissue (**Figure 4A and 4B**)^10^. Notably, multiple immune-related pathways were significantly downregulated in PB CRISPR tumors relative to normal muscle, including: B cell receptor signaling pathways, B cell activation pathways, activation of immune response pathways, and MHC class II protein complex assembly pathways (**Figure 4B**). These findings suggest that *Bap1* loss is associated with an immunosuppressive tumor microenvironment, consistent with previous observations in *BAP1*-driven uveal melanoma ^41^. Additionally, recent studies have shown that BAP1 loss leads to downregulation of MHC class II expression in B cell lymphoma ^26^.

**Figure 4.**
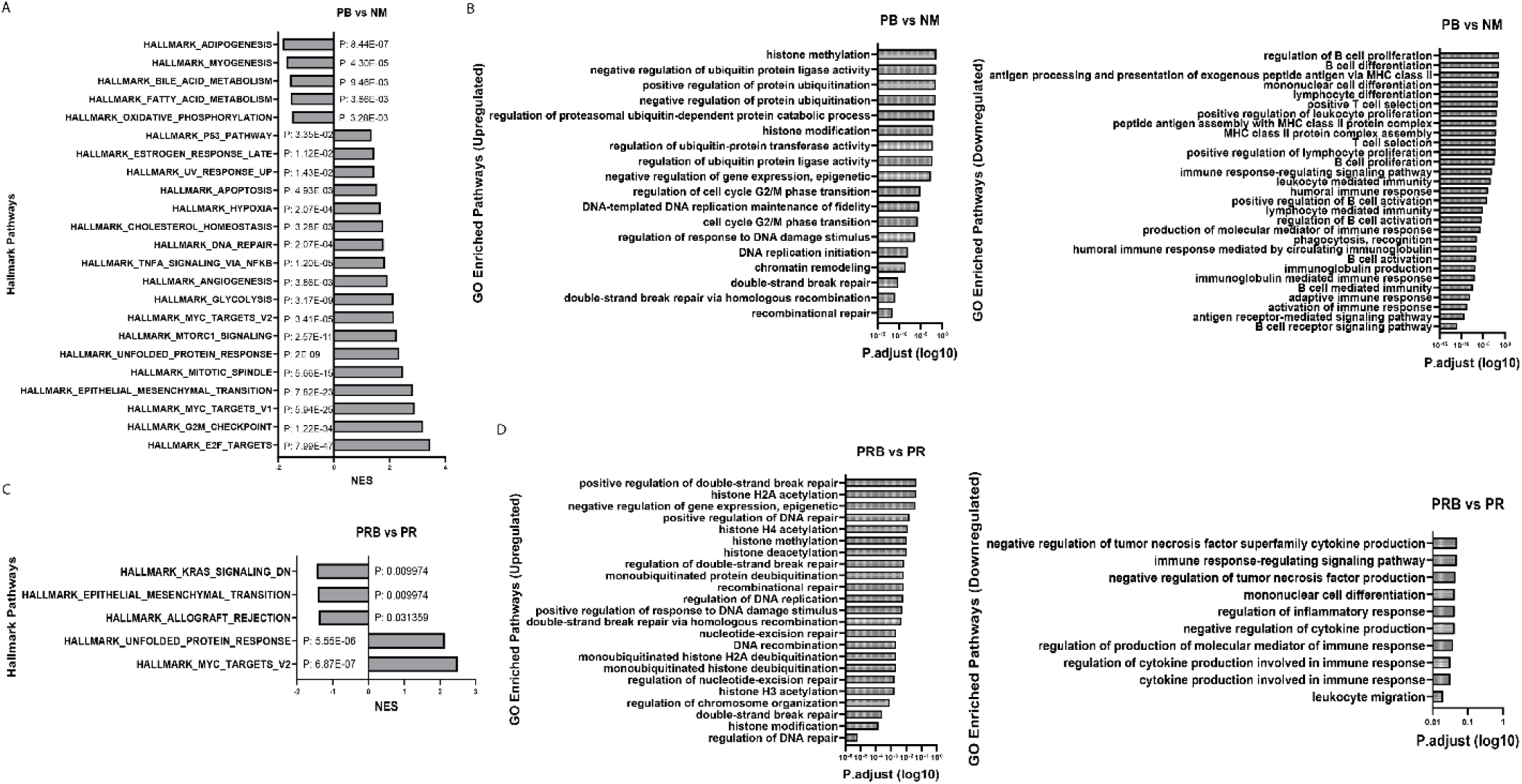
RNA sequencing reveals immune suppression in *Bap*-deficient sarcomas. (A-B) PB CRISPR tumors show upregulation of E2F targets, G2M checkpoint, MYC targets, DNA repair, and histone modification pathways, and downregulation of immune response pathways compared to normal muscle. (C-D) PRB Cre/CRISPR tumors exhibit upregulation of KRAS and MYC pathways and downregulation of B cell signaling and immune activation pathways compared to PR Cre tumors.

Given that *Bap1* deletion increased sarcoma aggressiveness in the context of *Trp53* and *Rb1* loss (**Figure 3D**), we next compared PR Cre tumors with PRB Cre/CRISPR tumors. PRB Cre/CRISPR tumors exhibited significant upregulation of oncogenic pathways, including: the hallmark of KRAS signaling, unfolded protein response and the hallmark of MYC targets (**Figure 4C**). These dysregulated pathways may contribute to the enhanced tumor penetrance and aggressiveness observed in PRB Cre/CRISPR tumors ^42,43^. Furthermore, PRB Cre/CRISPR tumors showed significant downregulation of immune-related pathways compared to PR Cre tumors, including: regulation of tumor necrosis factor (TNF) production, regulation of inflammatory response, and cytokine production in immune response (**Figure 4B**). Collectively, these results indicate that Bap1 loss contributes to immune suppression in soft tissue sarcomas, potentially facilitating tumor progression and resistance to immune surveillance.

### Immune cell profiles confirm an immunosuppressive microenvironment in *Bap1*-loss mouse sarcomas

To examine the tumor immune microenvironment (TIME) in mouse sarcomas driven by different genetic mutations, we performed multiplex immunohistochemistry (mIHC) ^44^ and flow cytometry (FC) on primary tumors ^45^. Using an mIHC panel including CD3, CD8, CD68, PD-1, CD20, and DAPI, we observed minimal infiltration of CD3⁺ T cells, CD8⁺ T cells, and CD20⁺ B cells in KP mouse sarcomas (driven by *Kras* ^G12D^ activation and *Trp53* deletion in *Kras* ^LSL-G12D/+^*; Trp53* ^FL/FL^ mice), consistent with previous reports (n = 1, data not shown) ^46,47^. In contrast, PR CRISPR tumors (n = 4) exhibited significantly higher infiltration of CD3⁺ T cells, CD8⁺ T cells, and CD20⁺ B cells compared to PB CRISPR tumors (n = 5) (**Figure 5A**). To further validate the impact of *Bap1* loss on immune cell infiltration, we compared PR Cre tumors (n = 3) with PRB Cre/CRISPR tumors (n = 4). PR Cre tumors showed significantly more CD3⁺ T cells, while CD8⁺ T cell and CD20⁺ B cell levels were comparable between groups (**Figure 5**). CD68⁺ macrophages were abundant across all tumor types, consistent with findings in human UPS ^48,49^ (**Supplementary Figure 8**).

**Figure 5.**
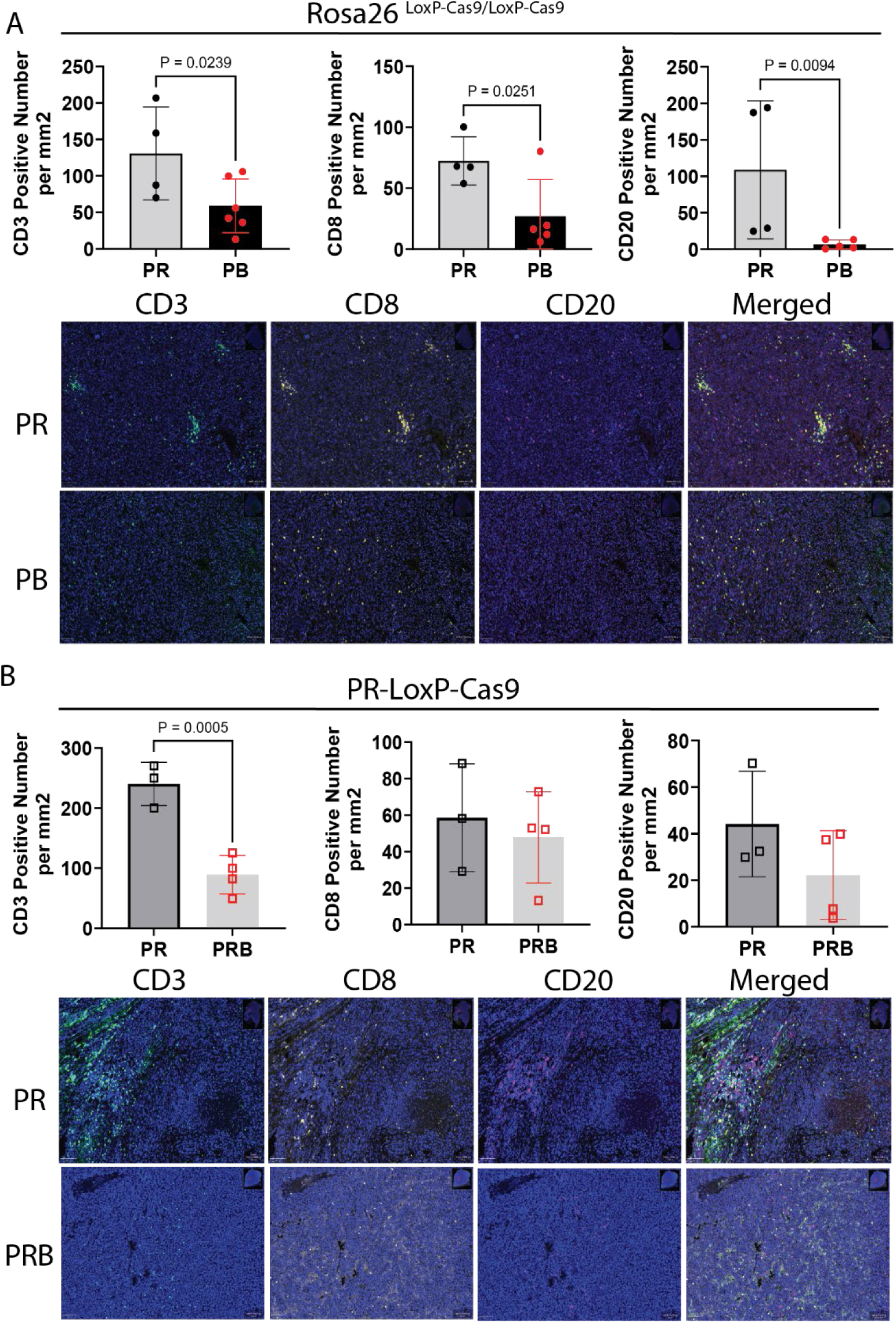
Multiplex immunohistochemistry reveals reduced immune cell infiltration in *Bap1*-deficient sarcomas. (A) PR CRISPR tumors (n = 4) show significantly more CD3⁺ T cells than PB CRISPR tumors (n = 6). (B) PR Cre tumors (n = 3) show significantly more CD3⁺ T cells than PRB Cre/CRISPR tumors (n = 4). Each point represents an individual tumor. Scale bar = 100 µm.

We next used FC to analyze immune cell populations in tumors of varying sizes. We compared normal muscle (NM; n = 5), PR CRISPR large tumors (n = 4), PB CRISPR small tumors (<250 mm³; n = 3), and PB CRISPR large tumors (>1000 mm³; n = 9) (**Figure 6A**). No significant differences were observed in CD8⁺ T cells among CD45⁺ cells. However, CD4⁺ T cell percentages were significantly lower in PB CRISPR large tumors compared to NM and PB early tumors. Notably, the ratio of CD4⁺ regulatory T cells (Tregs) to CD4⁺ T cells was significantly elevated in PB CRISPR large tumors. Further analysis revealed a higher frequency of proliferating (Ki-67⁺) CD8⁺ T cells in PR CRISPR large tumors compared to NM, PB large, and PB small tumors. Additionally, Ki-67⁺ Foxp3⁺ CD4⁺ Tregs were more abundant in PR CRISPR large tumors than in PB CRISPR large tumors. Although Ki-67⁺ Treg levels were lower in PB CRISPR large tumors than in PB small tumors, the difference was not statistically significant. PD-1 expression on CD8⁺ T cells did not differ significantly across tumor types (**Supplementary Figure 9**). These findings suggest that CD8⁺ T cells are more proliferative in PR CRISPR tumors than in PB CRISPR tumors.

**Figure 6.**
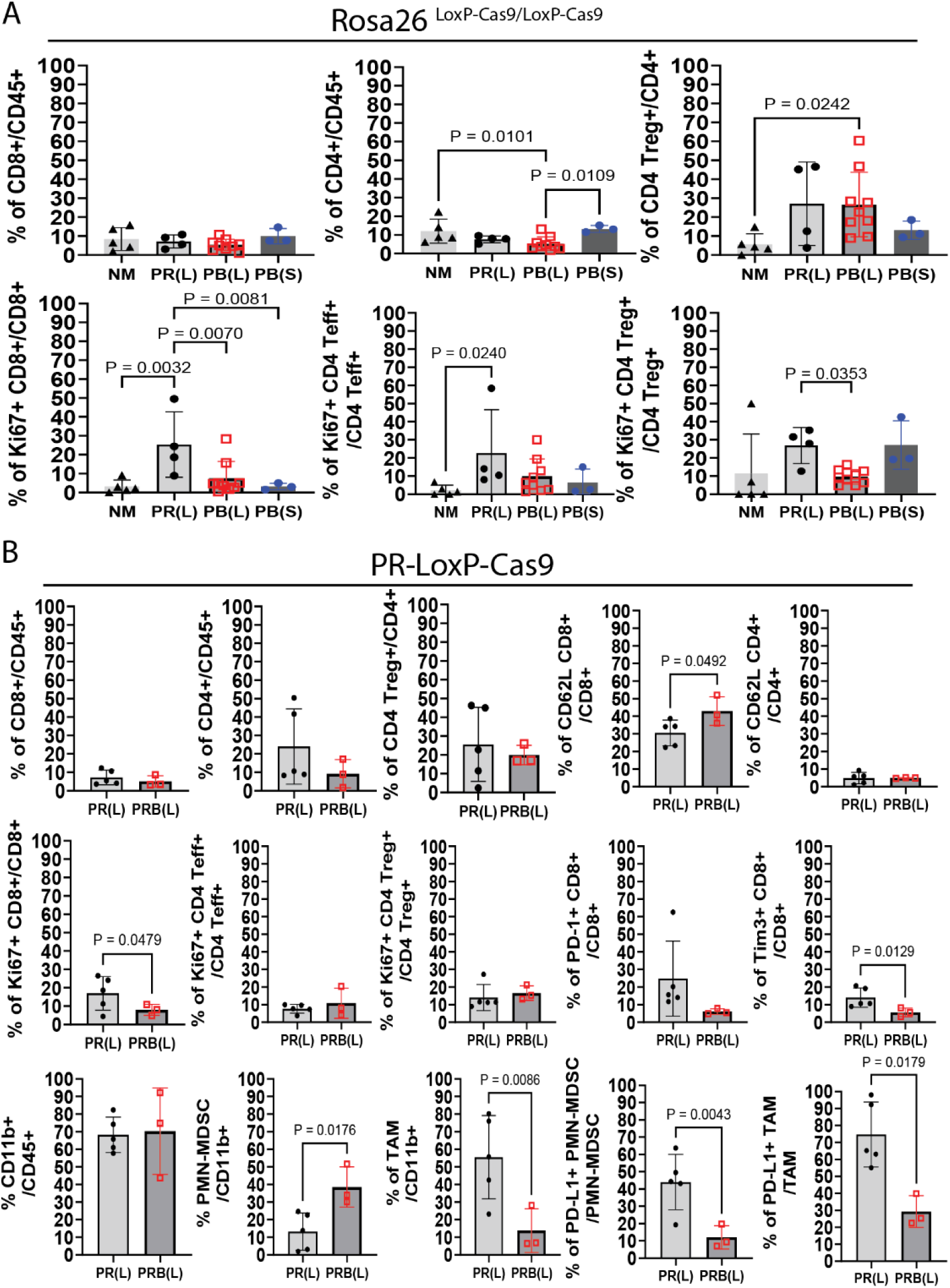
Flow cytometry analysis reveals immunosuppressive features in *Bap1*-deficient sarcomas. (A) No differences in CD8⁺ T cell percentages among CD45⁺ cells across NM (n = 5), PR CRISPR large (n = 5), PB CRISPR large (n = 9), and PB CRISPR small (n = 3) tumors. CD4⁺ T cell percentages were lower in PB CRISPR large tumors, with higher Treg/CD4⁺ ratios. PR CRISPR large tumors had more Ki-67⁺ CD8⁺ T cells and FoxP3⁻ CD4⁺ Teff cells. (B) No differences in CD8⁺, CD4⁺, or Treg cell percentages between PR Cre (n = 5) and PRB Cre/CRISPR (n = 3) tumors. PR Cre tumors had more Ki-67⁺ CD8⁺ T cells, Tim3⁺ CD8⁺ T cells, and CD11b⁺ cells. PRB Cre/CRISPR tumors had higher PMN-MDSC percentages and lower TAM percentages. PD-L1⁺ PMN-MDSCs were more abundant in PR Cre tumors. Each point represents an individual tumor or muscle.

We also profiled immune cells in PR Cre tumors (n = 5) and PRB Cre/CRISPR tumors (n = 3) (**Figure 6B**). PR Cre tumors exhibited significantly higher frequencies of proliferating (Ki-67⁺) CD8⁺ T cells and reduced expression of CD62L, a marker of T cell activation ^50^. Tim3⁺ CD8⁺ T cells were significantly lower in PR Cre tumors, while PD-1⁺ CD8⁺ T cell levels were similar between groups. These results suggest that CD8⁺ T cells in Bap1-wildtype tumors (PR Cre) are more proliferative and activated than those in Bap1-deficient tumors (PRB Cre/CRISPR).

We further examined myeloid-derived suppressor cells (MDSCs) and tumor-associated macrophages (TAMs). CD11b⁺ cell percentages among CD45⁺ cells were significantly higher in PB CRISPR large tumors compared to PB early tumors (**Supplementary Figure 10**). No differences were observed in CD11b⁺ cell percentages between PR Cre and PRB Cre/CRISPR tumors (**Figure 6B**). Additionally, there were no significant differences in polymorphonuclear MDSCs (PMN-MDSC; CD11b^+^-Ly6C^int^ -Ly6G^hi^), monocytic MDSCs (M-MDSCs; CD11b^+^-Ly6C^hi^-Ly6G^lo^), or TAMs (CD11b^+^-Ly6C^lo^-Ly6G^lo^-F4/80^+^) between PB CRISPR large and small tumors (**Supplementary Figure 10**). Interestingly, PMN-MDSC percentages were significantly higher in PRB Cre/CRISPR tumors, while TAM percentages were higher in PR Cre tumors. M-MDSC levels were similar between groups. PD-L1, an immunosuppressive ligand that binds PD-1 ^51^, was significantly lower on PMN-MDSCs, M-MDSCs, and TAMs in PRB Cre/CRISPR tumors compared to PR Cre tumors (**Figure 6B, Supplementary Figure 10**). Although PD-L1 expression is generally associated with poor prognosis in STS, its role in Bap1-deficient sarcomas warrants further investigation. ^52^. Overall, these findings confirm that Bap1 loss is associated with an immunosuppressive tumor microenvironment in STS, characterized by reduced T cell activation and proliferation, increased Treg ratios, and altered myeloid cell profiles.

### *Plk1* Knockdown inhibits growth of *Bap1*-loss mouse sarcomas

We established multiple mouse sarcoma cell lines derived from PR and PB CRISPR tumors and confirmed knockout of the targeted genes via western blotting (**Supplementary Figure 11**). Given prior evidence that PARP inhibitors (PARPi) are effective in *Bap1*-mutant renal cell carcinoma (RCC) ^53^, and that double-strand DNA repair pathways are upregulated in PB CRISPR tumors, we tested whether PARPi could eliminate PR and PB sarcoma cells. Additionally, PARPi have shown efficacy in *Rb1*-mutant osteosarcoma ^54^. Therefore, we tested whether PARPi could eliminate PR and PB sarcoma cells. In vitro live-cell imaging assays revealed that both PR and PB sarcoma cells were sensitive to two PARPi: niraparib and talazoparib (**Supplementary Figure 12**). However, in vivo treatment of nude mice bearing PR or PB tumors with niraparib (50 mg/kg daily via oral gavage for 28 days) showed significant anti-tumor effects only in PR tumors, not PB tumors (**Supplementary Figure 12**). These results suggest that PR tumors are sensitive to PARP inhibition in vivo, while PB tumors are resistant—consistent with clinical trial data showing limited efficacy of PARPi in *BAP1*-driven cancers ^55–57^.

Given the observed dysregulation of mitosis and chromosome segregation in PB sarcomas, we hypothesized that Polo-like kinase 1 (PLK1), a key regulator of mitosis and chromosome segregation, could be a therapeutic target ^29,30^. Using BAP1 chromatin immunoprecipitation followed by sequencing (ChIP-seq), we identified BAP1 binding at the PLK1 genomic locus (**Figure 8A**) ^26^. BAP1, a deubiquitylase, forms the PR-DUB complex to reverse H2AK119ub1 modifications catalyzed by PRC1 ^24,58^. *Bap1* knockout increased H2AK119ub1 levels upstream of the *PLK1* transcription start site (**Figure 8A**), suggesting epigenetic regulation of *PLK1* by BAP1 ^26^. Analysis of TCGA data revealed that high *PLK1* expression correlates with poor prognosis in UPS patients (**Figure 8B**). Total RNA sequencing and RT-PCR confirmed that *Plk1* is significantly upregulated in PB sarcomas compared to normal muscle (**Figure 8C**). To investigate the oncogenic role of PLK1, we stably transduced Cas9-expressing PB sarcoma cell lines with lentiviruses expressing either a non-targeting control sgRNA (neg sgRNA) or sgRNAs targeting *Plk1*. Most cells transduced with *Plk1* sgRNAs died, while control cells survived, despite equal viral titers (data not shown). We expanded the remaining *Plk1*-knockdown cells and confirmed reduced PLK1 expression via western blot (**Figure 8D**). In vitro proliferation assays showed that PLK1 knockdown significantly inhibited PB sarcoma cell growth (**Figure 8D**). To assess the effect of PLK1 knockdown in vivo, we injected Rosa26 ^LoxP-Cas9/LoxP-Cas9^ mice intramuscularly with PB sarcoma cells stably expressing neg sgRNA (n = 3), *Plk1* sgRNA1 (n = 4), or *Plk1* sgRNA2 (n = 3). PLK1 knockdown significantly suppressed tumor growth (**Figure 8E**). Notably, 2 out of 4 mice injected with *Plk1* sgRNA1-transduced cells did not develop tumors over a 72-day period. Moreover, PLK1 knockdown significantly prolonged mouse survival (**Figure 8E**). These findings suggest that PLK1 is critical for the growth of Bap1-deficient sarcomas and may represent a promising therapeutic target.

**Figure 7.**
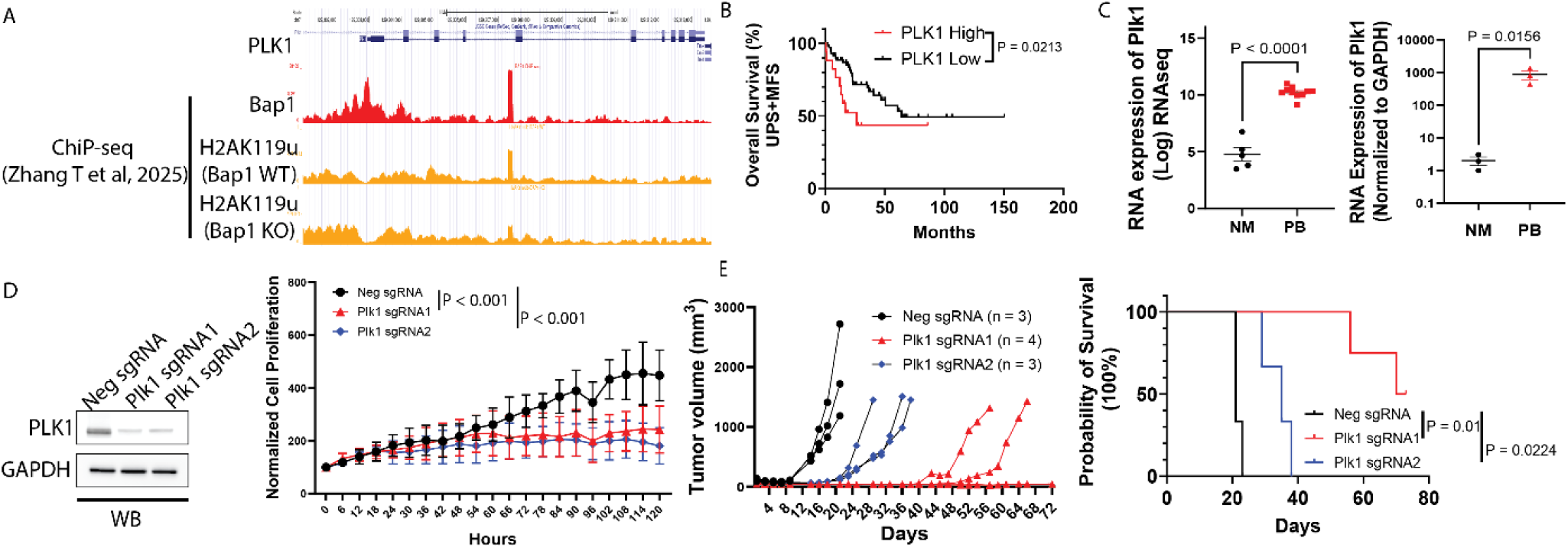
PLK1 Knockdown inhibits growth of *Bap1*-deficient sarcomas. (A) Bap1 binds the *Plk1* locus; Bap1 knockout (KO) increases H2AK119ub1 at the *Plk1* promoter. (B) High PLK1 expression correlates with poor prognosis in UPS (TCGA). (C) RNA-seq and RT-qPCR confirm *Plk1* upregulation in PB sarcomas. (D) Western blot confirms PLK1 knockdown; Cellcyte assay shows reduced proliferation. (E) PLK1 knockdown inhibits tumor growth and prolongs survival in syngeneic mouse model.

**Figure 8.**
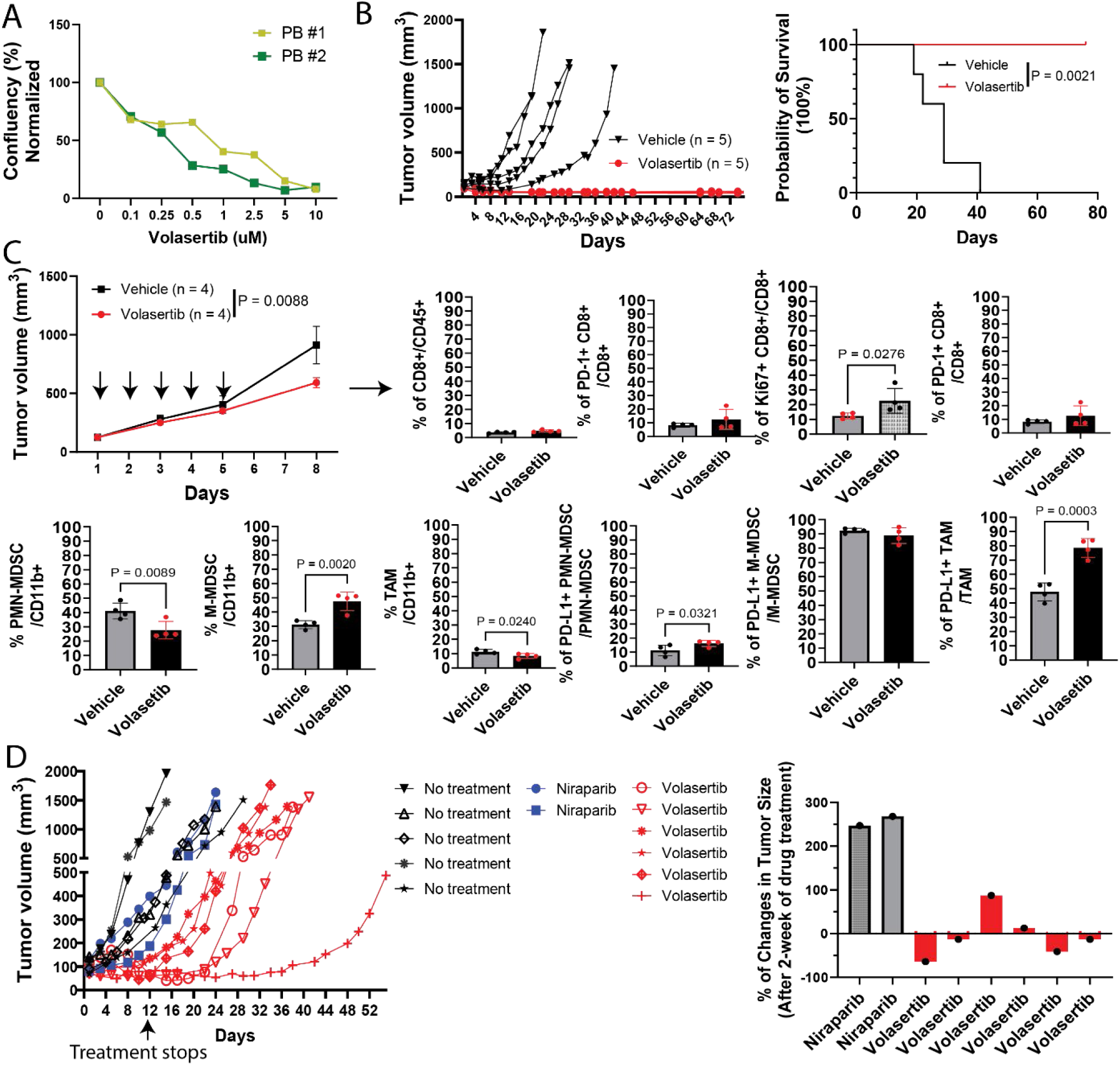
Pharmacological inhibition of PLK1 suppresses Bap1-deficient sarcomas. (A) Volasertib inhibits PB sarcoma cell proliferation in vitro in a dose-dependent manner. (B) Volasertib suppresses tumor growth and prolongs survival in B6 mice bearing PB sarcomas. (C) In Rosa26 ^LoxP-Cas9/LoxP-Cas9^ mice, volasertib increases Ki-67⁺ CD8⁺ T cells, reduces TAMs, and alters MDSC populations. PD-L1⁺ PMN-MDSCs and TAMs are elevated post-treatment. (D) In autochthonous UPS model, volasertib induces tumor regression in 4/6 mice, while niraparib shows no effect. Left: tumor growth curves; right: waterfall plot of tumor size changes after 2-week treatment.

### Pharmaceutical inhibition of Plk1 as a targeted therapy for *Bap1*-loss sarcomas

To evaluate the therapeutic potential of PLK1 inhibition in *Bap1*-deficient sarcomas, we tested the sensitivity of PB mouse sarcoma cells to volasertib, a selective PLK1 inhibitor ^59^. In vitro assays demonstrated that PB sarcoma cells were sensitive to volasertib in a dose-dependent manner (**Figure 9A**). We next assessed volasertib’s efficacy in vivo using immune-competent C57BL/6J (B6) mice transplanted intramuscularly with PB sarcoma cells. Treatment began when tumors reached 50–100 mm³, with mice receiving 10 mg/kg volasertib or vehicle via oral gavage, five days per week for two weeks. Volasertib significantly suppressed tumor growth (**Figure 9B**), and tumors remained undetectable in all five treated mice up to 60 days post-treatment. These results indicate that volasertib prolongs survival and induces durable tumor regression in PB sarcoma-bearing mice. To investigate volasertib’s impact on tumor-infiltrating immune cells, we used Rosa26 ^LoxP-Cas9/LoxP-Cas9^ mice injected intramuscularly with PB sarcoma cells. Treatment began when tumors reached ≥100 mm³, with mice randomized to receive volasertib or vehicle (10 mg/kg, 5 days/week for 1 week) (**Figure 9C**). Flow cytometry performed on Day 4 post-treatment revealed: Significantly increased frequencies of proliferating (Ki-67⁺) CD8⁺ T cells in the volasertib group; No difference in CD11b⁺ cell percentages among CD45⁺ cells; Significantly lower percentages of PMN-MDSCs and TAMs, but higher percentages of M-MDSCs in the volasertib group; Higher expression of PD-L1 on PMN-MDSCs and TAMs in volasertib-treated tumors (**Supplementary Figure 13**, **Figure 9C**). These findings suggest that volasertib not only suppresses tumor growth but also modulates the immune microenvironment, enhancing CD8⁺ T cell proliferation and altering myeloid cell composition.

Finally, we tested volasertib in spontaneous tumors initiated by IVE delivery of *Trp53* and *Bap1* sgRNAs into the gastrocnemius muscle of Rosa26 ^LoxP-Cas9/LoxP-Cas9^ mice (**Figure 3A**). When tumors reached 50–100 mm³ (≥100 days post-injection), mice were treated with either 50 mg/kg niraparib (n = 2) or 10 mg/kg volasertib (n = 6) for two weeks. Niraparib showed no therapeutic effect, consistent with transplantation model results. In contrast, volasertib induced tumor regression in 4 out of 6 mice, with one tumor remaining stable and one showing slow growth (**Figure 9D, waterfall plot**). Overall, these results demonstrate that pharmaceutical inhibition of PLK1 via volasertib is a promising targeted therapy for Bap1-loss sarcomas, with both direct anti-tumor effects and beneficial modulation of the tumor immune microenvironment.

## Discussion

Undifferentiated pleomorphic sarcoma (UPS), a major subtype of soft tissue sarcoma (STS) in adults, is associated with high mortality due to recurrence and metastasis, with a median survival of less than 15 months ^2,3,7^. Despite its clinical severity, treatment options for UPS have remained largely unchanged for over four decades. The genetic heterogeneity and absence of recurrent oncogenic driver mutations in UPS make a universal therapeutic approach impractical. Instead, UPS frequently exhibits copy number alterations in tumor suppressor genes, suggesting that identifying actionable targets based on specific tumor suppressor mutations may offer a more effective strategy for therapy development.

A deeper understanding of the mechanisms driving UPS development and immune evasion is urgently needed. However, the lack of spatially and temporally controlled autochthonous mouse models that accurately mimic human UPS has hindered progress. Our previous in vitro genome-scale CRISPR/Cas9 screen failed to recapitulate sarcomagenesis in vivo ^37^. To overcome this limitation, we developed a tailored in vivo CRISPR/Cas9 screen in mice, identifying tumor suppressor genes whose mutation, in combination with *Trp53* loss, drives UPS formation. This approach revealed that additional mutations beyond *Trp53* are required for sarcoma initiation.

Through individual gene validation, we demonstrated that concurrent mutations in *Bap1* and *Trp53* are sufficient to induce sarcomagenesis. While *BAP1* mutations are present in a small subset of STS, they are predominantly found in UPS according to TCGA data ^60^. Moreover, *BAP1* mutations have been reported in 12% of recurrent pediatric embryonic rhabdomyosarcomas, which share a similar cellular origin with UPS ^61–64^, and in patients with BAP1 tumor predisposition syndrome (BAP1-TPDS) ^18,65,66^. To our knowledge, this study provides the first direct evidence that *Bap1* loss contributes to sarcoma initiation, challenging the prevailing view that *BAP1* mutations are late events in tumor progression ^23^. Our UPS model driven by somatic deletion of *Trp53* and *Bap1* offers a unique platform to investigate BAP1’s role in tumorigenesis and therapeutic targeting.

*BAP1* functions as a deubiquitylase within the polycomb repressive-deubiquitylase (PR-DUB) complex, which includes Host cell factor 1 (*HCFC1*), O-linked N-acetylglucosamine transferase (*OGT*), Forkhead box K1/2 (*FOXK1/2*) and Additional sex combs like 1/2/3 (*ASXL1/2/3*) ^28^. Mutations in *ASXL1* are common in hematologic malignancies, while *ASXL2* and *ASXL3* are implicated in other cancers ^67^. *FOXK1* and *FOXK2* also play critical roles in cancer development ^68^. Notably, TCGA database show that *FOXK1*, *FOXK2*, *ASXL2*, *ASXL3*, and *BAP1* are exclusively mutated or copy number altered in UPS (**Supplementary Figure 14**). A total of 16% of human UPS bear mutations or deletions in at least one gene from the PR-DUB complex. A recent study using CRISPR/Cas-mediated saturation genome editing screening determined that the *ASXL1/2/3* interacting domain in the BAP1 protein may be critical for its tumor suppressive function ^69^. These findings suggest that disruption of the PR-DUB complex by a single gene mutation may promote sarcoma development, an observation that warrants further study.

Beyond tumorigenesis, *BAP1* also influences the tumor immune microenvironment. In uveal melanoma, loss of *BAP1* is significantly correlated with an immunosuppressive microenvironment ^41,70^. *Bap1* deletion resulted in resistance to PD-1 blockade in a synergic mouse models of pancreatic cancer driven by activation of *Kras* ^G12D^ and deletion of *Trp53*. In our study, RNA sequencing revealed significant downregulation of B cell and immune response pathways in *Bap1*-deficient sarcomas compared to controls. Additionally, mIHC and FC confirmed poor immune cell infiltration, particularly of proliferating CD8^+^ T cells. Prior research has shown that ablation of BAP1 reduces transcription of major histocompatibility complex class II (MHC-II) in B cell lymphomas, contributing to immune suppression ^26^. Our RNA sequencing result similarly show MHC-II downregulation in Bap1-deficient, though its role in STS immune evasion remains to be elucidated (data not shown). Future studies will focus on dissecting how Bap1 regulates the tumor immune microenvironment in STS.

Targeted therapy holds promise for genetically diverse UPS. For instance, Li G et al. showed that *RB1* and *TP53*-deficient UPS and MFS are sensitive to SKP2 inhibition. Studies showed that *RB1*-mutant osteosarcomas and *BAP1*-mutant RCC are sensitive to PARPi ^53,54^. We investigated the effect of PARPi on PR and PB mouse sarcomas. We found that all tested mouse sarcoma cells showed similar sensitivity to two PARPi, niraparib and talazoparib. However, only PR sarcomas showed significant in vivo sensitivity to niraparib, while PB sarcomas were resistant, which is consistent with clinical observations of *BAP1*-mutant tumors being refractory to PARPi ^55–57,71^. Therefore, we explored alternative actionable targets such as Plk1, a serine/threonine protein kinase that plays a critical role in tumor progression, survival, immune suppression ^31–33,72,73^. Upregulation of *PLK1* is associated with poor prognosis of human UPS. We showed that *Plk1* is critical for the survival of *Bap1*-deficient sarcomas. Utilizing both transplantation and spontaneous mouse models, we demonstrated that *Bap1*-deficient sarcomas are sensitive to the PLK1 inhibitor, volasertib. Furthermore, volasertib treatment had a positive impact on tumor infiltrating immune cells in *Bap1*-deficient sarcomas. These findings suggest that *Plk1* inhibition may be a viable therapeutic strategy and could potentially be combined with immunotherapy in *BAP1*-deficient UPS.

In conclusion, this study establishes a novel autochthonous mouse model of UPS driven by *Bap1* and *Trp53* loss, providing critical insights into the genetic and immunological mechanisms of sarcomagenesis. Our findings challenge the notion that BAP1 mutations are late events and highlight their role in tumor initiation and immune evasion. The identification of Plk1 as a therapeutic vulnerability in Bap1-deficient sarcomas opens new avenues for targeted treatment, particularly in combination with immunotherapy. This model lays the groundwork for future studies aimed at unraveling the complex interplay between tumor suppressor mutations and the immune microenvironment in UPS.

## Methods

### Mice

Rosa26 ^LoxP-Cas9^ mice were provided by F. Zhang (Massachusetts Institute of Technology). Trp53 ^Flox^ mice were provided by A Berns (University of Amsterdam). Rb1 ^Flox^ mice were provided by I Gelman (Roswell Park Cancer Institute). Fat1 ^Flox^ mice were provided by N Sibinga (Albert Einstein College of Medicine). For the transplantation study, male athymic nude (nu/nu) mice (6 weeks old) were purchased from the Jackson Laboratory. The nude mice were maintained in Earle A. Chiles Research institute’s accredited animal facility. Both male and female mice were used in experiments. All animal studies were performed in accordance with protocols approved by the Animal Care and Use Committee of the Earle A. Chiles Research Institute.

### Cell culture

Mouse embryonic fibroblasts were generated from E13.5 to E14.5 embryos using standard procedures. NIH-3T3 cell lines (CRL-1658) were purchased from ATCC. They were cultured in DMEM medium (Cytiva, SH30243.LS) supplemented with 10%fetal bovine serum and 1% penicillin-stregtomycin and L-Glutamine (Corning, 30-009-CI). Cells were incubated at 37 °C with 5% CO_2_ in a cell-culture incubator. Mouse sarcoma cells were isolated and maintained in cell culture from primary tumors following standard methods ^3^. Tumor tissues were minced, and then digested by dissociation buffer containing collagenase type IV (Thermo Fisher Scientific, 17104-019), dispase (Thermo Fisher Scientific, 17105-041) and trypsin (Thermo Fisher Scientific, 25200056) for 1 h in a shaker at 37 °C. Cells were washed with 1 × PBS (Thermo Fisher Scientific, SH30256LS) and filtered with a 40 mm sieve (Corning, 43175).

### Plasmids

The Cre recombinase in the px333-Cre vector was deleted using two NcoI sites to generate px334 vector by standard cloning methods. For cloning px334-Negative sgRNA, px334-Trp53 sgRNA, and px334-Cdkn2a sgRNA, the px334 vector was digested with BsaI enzyme and ligated to annealed sgRNA oligonucleotides (**Supplementary Table 2**). For cloning customized sgRNA libraries into these three vectors, a 35-gene targeting sgRNA library plus control sgRNAs were taken from the mouse Brie library ^36^. Oligonucleotide pools (**Supplementary Table 5**) were ordered from Twist Bioscience and cloned into the three px334 vectors by Gibson assembly (New England Biolabs, E2611), as previously described ^37^. In brief, transformations of the assembled library were done into Endura competent cells (Lucigen, 60242-1) and sgRNA representation was verified by high-throughput sequencing. Additional 3 genes (Ano7, Csmd1, and Myt1l) that are frequently mutation in myxofibrosarcoma were not included in oligonucleotide pools. Therefore, gene targeting sgRNAs were cloned into px334 vector, and those vectors were proportionally mixed with the library plasmids before injection into the mice (**Supplementary Table 4**). For cloning individual candidate gene targeting sgRNAs into the px334 or px333-Cre vectors, the vector was digested with BbsI enzyme and ligated to annealed sgRNA oligonucleotides (**Supplementary Table 4**).

### Tumor analysis

Mouse tissues were harvested, fixed in 4% formalin, and embedded in paraffin. Immunohistochemistry was performed on 5 µm sections of mouse tissues with the ABC kit (Vector Laboratories, PK-7200) with antibodies to MyoD (Agilent Technologies, M351201-2), Myogenin (Agilent Technologies, IR06761-2), Cytokeratin (Agilent Technologies, GA0536-2), SMA (Agilent Technologies, GA61161-2), Desmin (Agilent Technologies, GA63061-2), S100 (Agilent Technologies, GA50461-2), Yap1 (Cell Signaling Technology, 4912S). Tissue sections were developed with 3,30 -diaminobenzidine and counterstained with hematoxylin (Sigma-Aldrich, H3136). Hematoxylin and eosin staining was performed using standard methods. All tissue sections were blindly examined by a sarcoma pathologist.

### Genomic DNA isolation and indel analysis

Genomic DNA was isolated from both cells and tissues using Zymo Quick-DNA Miniprep kit (Zymo Research, D3024). PCR amplification of mouse targeted genes by specific sgRNAs was performed with specific primer (**Supplementary Table 4**) using Q5 HiFi DNA polymerase (New England Biolabs, M0494). PCR reaction was sent to Azenta for Sanger sequencing or Next Generation Sequencing.

### Western blot analysis

NIH-3T3 and mouse sarcoma cell lines were treated with 0.5 µg/ml doxorubicin (Sigma-Aldrich, D1515-10MG) for 16 h before sample harvest. Samples were lysed in RIPA buffer containing protease inhibitor cocktails (Sigma-Aldrich, R0278) for 30 min on ice, then centrifuged at 15,000 for 2 min. Lysate supernatant was separated from debris into a new tube and protein concentration was determined by Nanodrop. The lysate was boiled in 2 × sample buffer (Thermo Fisher Scientific, 84788) at 100 °C for 5 min, then cooled to room temperature before loading in a 4-20% Mini-Protean TGX precast protein gels (Bio-Rad, 4561093). Samples were electrophoresed at 100 V for 60 min before transfer to PVDF (Thermo Fisher Scientific, IB24002) using Iblot 2 Dry Blotting System (Thermo Fisher Scientific, IB21001). The membrane was blocked in 5% non-fat dry milk in tris-buffered saline for 1 h (TBS, Corning, 46-012-CM). Next, membranes were incubated with primary antibodies diluted in TBS-T (0.1% Tween-20) in a shaker overnight at 4 °C: p53, 1:1000 dilution (Cell Signaling Technology, 2524s); Rb1, 1:500 dilution (Santa Cruz, sc-102), Bap1, 1:1000 dilution (Cell Signaling Technology, 13271s), and Gapdh, 1:100000 dilution (Proteintech, 60004-1-lg). Membranes were washed three times in TBS-T for 5 min before incubation with Pierce HRP Goat anti-rabbit secondary antibody (Fisher Scientific, PI31460) and Pierce HRP Goat anti-mouse secondary antibody (Fisher Scientific, PI31430) both at 1:10,000 dilutions in TBS-T for 1 h at room temperature. The membranes were washed three times in TBS-T for 5 min and imaged using a Bio-Rad ChemiDoc imaging system. Full blots with protein ladders (Li-Cor Biosciences, P/N 928-60000) are shown in **Supplementary Figure 15**.

### Multiplex immunohistochemistry

Mouse FFPE tissue slides were dewaxed in xylene and rehydrated in decreasing graded alcohol followed by sequential stain. The staining procedure was performed as the protocol published from EACRI IHC Core Lab ^44^. Antigen retrieval was done in Tris-EDTA, pH9.0 in microwave oven. Citrate buffer, pH6.0 was used for antibody stripping. 3% H_2_O_2_ was used for killing exogenous HRP when followed biomarker has antibody raised from different species of animal. Tissue sections were blocked with blocking/antibody diluent (ARD1001EA, Akoya) for 10 min at room temperature (RT). Anti-CD68 (1:1200, E3O7V, Cell Signaling), anti-PD-1(1:200, D7D5W, Cell Signaling), anti-CD8 (1:400, 4SM15, eBioscience), anti-CD3(1:100, SP7, Genetex), and anti-CD20(1:6400, E3N7O, Cell Signaling) were sequentially stained. After a brief wash, tissue sections were incubated with MACH2 Rb HRP-Polymer (RHRP520H, Biocare Medical) or Rat HRP-Polymer, (BRR4016 H, Biocare Medical) for 10 min at RT. After a quick wash, Opal690(1:200, Akoya), Opal620 (1:200, Akoya), Opal570 (1:400, Akoya), and Opal520 (1:200, Akoya), respectively, for 10 min at RT, Opal780 (1:25, Akoya) for 60 min at RT. DAPI stain (1 drop of DAPI solution into 0.5ml of TBST, Akoya) was done in 5 min at RT. Then the slides were mounted with Prolong Diamond Antifade Mountant (p36970, Thermofisher). Imaging was performed on PhenoCycler-Fusion. Whole slide imaging was performed using a 20x objective lens, while high power multispectral images were done using a 40x objective lens. Images were analyzed using QuPath software ^74^.

### Flow cytometry

Mouse sarcoma tumors and normal muscle were harvested, chopped finely and collected directly into 2mL digest buffer for 20-45 minutes on a rocker/shaker. The digestion buffer was prepared with 0.25mg/ml Collagenase H (Sigma, Cat#C8051) and 30U/ml DNaseI (Roche, Cat#04536282001) in cDMEM. Single cell suspension was collected after filtering and washing the tumor digestions in 70 μM Tube top strainers (CellTreat, Cat#229483) with cDMEM. The single cell suspensions were then stained with combinations of surface markers and intracellular targets (**Supplementary Table 6**) following the method by Rolig AS et al (2022) ^39^. Flow cytometry data were acquired on Cytek Aurora flow cytometer (Cytek Biosciences) and data were processed and analyzed with OMIQ software (Dotmatics).

### Total RNA sequencing and analysis

Mouse sarcoma and normal muscle samples were preserved in RNAlater (Sigma-Aldrich, R0901-100ML). Then total RNA was isolated from samples using Quick-RNA Miniprep kit (Zymo Research, R1055). Then RNA was prepared with TruSeq Stranded Total RNA library prep kit with Ribo-Zero Human/Mouse/Rat Set Q (Illumina), and then sequenced with NovaSeq 6000 S1 Reagent Kit v1.5 at 100 cycles (Illumina, 20228318). Analysis of RNA-seq data is followed with a previous study ^3^. In brief, the quality of the sequencing reads was assessed using FastQC. The reads were aligned to the UCSC mm10 mouse genome from the iGenome Project using STAR and mapped to the mouse transcriptome annotated by GENCODE (Release M14). Raw gene counts were quantified using the HTSeq tool implemented in the STAR pipeline. Differentially expressed genes were first identified by modeling the raw counts within the framework of a negative binomial model using the R package DESeq2. Log2 fold change values were shrunk using the “apeglm” method. Gene set enrichment analysis was performed on mouse hallmark gene sets from MSigDB using the R package fgsea. Gene Ontology (GO) term enrichment was performed on biological process GO terms using the R package clusterProfiler. All P values were adjusted for multiple testing using the Benjamini–Hochberg method. The analyses were scripted in the R statistical environment (https://www.R-project.org/) along with its extension packages from the Comprehensive R Archive Network (https://cran.r-project.org/) and the Bioconductor Project. The raw data has been deposited in Sequence Read Archive (SRA) (PRJNA1212052).

### RNA extraction and real time qPCR

RNA was extracted from tissues using the Quick-RNA miniprep kit purchased from Zymo Research (R1058). cDNA was prepared using LunaScript RT SuperMix kit purchased from New England Biolabs (E3010L). RT-qPCR was performed in biological duplicates using QuantStudio 6 System (Applied Biosystems). RNA levels of Plk1 are quantified using a primer set (Forward: cagcagcaggaaacctctca; Reverse: caggatcctcagcctcctct).

### In vitro CellCyte Assay

Mouse sarcoma cell lines of the drug screening were cultured in complete DMEM (cDMEM) medium as described above. Cells were inoculated into 96-well microtiter plates in 100ml at plating densities ranging from 2,000 to 4,000 cells/well. After cell inoculation, the microtiter plates were incubated at 37° C, 5 % CO2, 95 % air and 100 % relative humidity for 24 h prior to addition of experimental drugs. The experimental drugs were solubilized in dimethyl sulfoxide (DMSO) at 400-fold or higher than the desired final maximum test concentration and then diluted to twice the desired test concentrations with cDMEM. DMSO was diluted with the volume in the desired maximum test concentration in cDMEM and used as the vehicle control. Aliquots of 100 μl of these different drug dilutions were added to the appropriate microtiter wells already containing 100 μl of medium, resulting in the required final drug concentrations. Following drug addition, the plates were loaded in The CELLCYTE X^TM^ live-cell imaging system (Cytena, Freiburg im Breisgau, Germany), incubated at 37°C, 5 % CO2, 95 % air, and 100 % relative humidity. The cells were monitored for 120 hours, two images per well were acquired every 6 hours using the 10X objective of the CELLCYTE X^TM^. A total of 21 scans were performed. The images were analyzed using the CELLCYTE^TM^ Analysis software to generate cell confluence data. The experiments were performed in duplicate.

### Chemical Compounds used in in vitro and in vivo study

Niraparib was purchased from TargetMol (T3231). Talazoparib was purchased from MedChem Express (HY-16106). Volasertib was purchased from AdooQ Bioscience (A10135) or TargetMol (T6019). For in vitro study, all compounds were dissolved in DMSO. For in vivo study, volasertib was resuspended in 0.5% methylcellulose.

### Targeted Capture Sequencing

The Amplicon targeted-capture sequencing probes were synthesized through Azenta using the method previously described ^34^. The capture sequencing was done in Azenta. The raw data has been deposited in Sequence Read Archive (SRA) (SUB15020645).

### Significantly mutated genes

Mutect2 and Varscan were used to call mutations. Mutect2 mutations must have at least 1000 reads, have a variant frequency of at least 5%. Varscan mutations must have a total of 100 reads and a variant frequency of at least 10%. Filtered mutations used in downstream analyses are Mutect2 mutations that are either flagged as “PASS” or are Varscan mutations as well. Furthermore, these filtered mutations must be located within the probe targeted regions. Annotations of Mutect2 mutations were done via SnpEFF.

### *In vivo* electroporation

All in vivo electroporation was performed with the method as previously described ^11^. Mice furs were shaved at least 1 day prior to the experiment. Then, 50 µg of endotoxin-free DNA plasmids in 50 µl diluted in sterile saline was intramuscularly injected into the mice using an 28g syringe. Next, a pair of needle electrodes was inserted into the muscle at the injection site, and electric pulses were delivered using a BTX ECM830 device (Harvard Apparatus, 45-2052). sgRNAs used in the individual gene validation experiments are all from the customized sgRNA library listed in the supplementary Table S2, except Trp53 sgRNA (gtgtaatagctcctgcatgg), Negative control sgRNA (GCGAGGTATTCGGCTCCGCG), Cdkn2a sgRNA (GGGCCGCCCACTCCAAGAGA), and Pten sgRNA (GCTAACGATCTCTTTGATGA). To mutate Trp53 and Rb1 in the Rosa26 ^LoxP-Cas9^ mice, we delivered a mixed plasmid pool of pX333-Trp53-Rb1 sgRNA1 (AAATGATACGAGGATTATCG), pX333-Trp53-Rb1 sgRNA2 (AGAGAAGTTTGCTAACGCTG), pX333-Trp53-Rb1 sgRNA3 (TAAGTACGTTCAGAATCCAC), and pX333-Trp53-Rb1 sgRNA4 (GCAGTATGGTTACCCTGGAG). We also generate tumors in mice with similar tumor onset by delivering a single plasmid of pX333-Trp53-Rb1 sgRNA4. To mutate Trp53 and Pten in the Rosa26 ^LoxP-Cas9^ mice, we delivered a plasmid of pX333-Trp53-Pten sgRNA. To mutate Trp53 and Bap1 in the Rosa26 ^LoxP-Cas9^ mice, we delivered a mixed plasmid pool of pX333-Trp53-Bap1 sgRNA1 (CCACCAACGTAGAAACCTTG) and pX333-Trp53-Bap1 sgRNA4 (TCAGCTATGTGCCTATCACA). We also generate tumors in mice with similar tumor onset by delivering a single plasmid of pX333-Trp53-Bap1 sgRNA4. To mutate Trp53 and Crebbp in the mice, we delivered a mixed plasmid pool of pX333-Trp53-Crebbp sgRNA3 (TAATGAATCAGGCTCAACAA) and pX333-Trp53-Crebbp sgRNA4 (TGAACCTACTGAATCCAAGG). To mutate Trp53 and Mst1r in the mice, we delivered a mixed plasmid pool of pX333-Trp53-Mst1r sgRNA1 (AAGTATCAGACTTTAGACGA) and pX333-Trp53-Mst1r sgRNA2 (GGGAACACACCAGATCACCG). To mutate Trp53 and Pbrm1 in the mice, we delivered a mixed plasmid pool of pX333-Trp53-Pbrm1 sgRNA1 (AAAACACTTGCATAACGATG) and pX333-Trp53-Pbrm1 sgRNA3 (AATAAAAGAGCAGTCCAAGG). To mutate Trp53 and Cysltr2 in the mice, we delivered a mixed plasmid pool of pX333-Trp53-Cylstr2 sgRNA1 (AAACCCATATGATCCCACAG) and pX333-Trp53-Cysltr2 sgRNA4 (GATGAATAGAAAATCGGAAG). To mutate Trp53 and Mllt3 in the mice, we delivered a mixed plasmid pool of pX333-Trp53-Mllt3 sgRNA3 (TCCACGATGTCATCAAACGG) and pX333-Trp53-Mllt3 sgRNA4 (ACTTACTCACCGTCACCAGT). To delete Trp53 and Rb1 in the Trp53 ^FL/FL^; Rb1 ^FL/FL^; Rosa26 ^LoxP-Cas9/LoxP-Cas9^ mice, we delivered the px333 plasmid to the mice. To delete Trp53, Rb1, and Bap1 in the Trp53 ^FL/FL^; Rb1 ^FL/FL^; Rosa26 ^LoxP-Cas9/LoxP-Cas9^ mice, we delivered a mixed plasmid pool of pX333-Trp53-Bap1 sgRNA1 and px333-Trp53-Bap1 sgRNA4.

### Statistical analysis

Results are presented as means ±SEM unless otherwise indicated. Before analysis, all data were displayed graphically to determine whether parametric or non-parametric tests should be used. Two-tailed Student’s t-test was performed to compare the means of two groups. To test the difference between groups, one-sided Wilcoxon rank-sum test was used for unpaired samples. To compare tumor-free survival, Kaplan-Meier analysis was performed with the log-rank test for statistical significance. Significance was assumed at P < 0.05. All calculations were performed using Prism 10 (GraphPad).

## Acknowledgements

We thank flow cytometry core and Philip D. Sanders for flow cytometry and technical support. We thank Grace H. McGee for technical support on flow cytometry. We also thank south wing lab meeting and attendees for valuable feedback on the manuscript. We also thank Allie Grossmann for valuable advice on the histology of mouse sarcomas. This work was supported by the National Cancer institute of the US NIH under award numbers 5 K22 CA248849 (J.H.), Sarcoma Foundation of America grant (J.H.), R35CA197616 (D.G.K.), and the Providence Portland Medical Foundation.

## Author contributions

J.H., and D.G.K. designed experiments. J.H., X.L., W.H., Z.S., N.T.W., M.J.K., W.F., R.P., S.Y.K., E.X., L.L., Y.M., Z.Z., and K.M.O. performed experiments. W.H. and Y.M. performed IHC. Z.S. performed mIHC. J.H., X.L., and M.J.K. perform FC. Y.W. examined histology of mouse sarcomas. B.P., J.T.W., V.R., W.K.R. and B.B. performed and analyses deep sequencing. R.B.B., W.L.R., W.J.U., provided critical advice for the manuscript. J.H., W.J.U., and D.G.K. drafted the first version of the manuscript. All authors edited the manuscript.

