## Supplementary figures and tables for "A tailored *in vivo* CRISPR screen identifies *BAP1* as a potent tumor suppressor of soft tissue sarcoma"

NC 27710

[6] Radiation Medicine Program, Princess Margaret Cancer Center, University Health Network, Toronto, ON, Canada

[7] Department of Radiation Oncology, University of Toronto, Toronto, ON, Canada

[8] Department of Medical Biophysics, University of Toronto, Toronto, ON, Canada

[#] Current position: Division of Radiation Oncology, University of Texas MD Anderson Cancer Center, Houston, TX 77030

[$] Department of Radiation Oncology, Baylor College of Medicine, Houston, TX 77030

[*] Corresponding authors: Jianguo Huang, Earle A. Chiles Research Institute, Providence Cancer Institute, Portland, OR 97213, and David G. Kirsch, Department of Radiation Oncology, Radiation Medicine Program, University of Toronto, Princess Margaret Cancer Center, Toronto, ON, Canada

**Supplementary Figure 1**. Mutation rates of three genes (TP53, Rb1, and BAP1) in human UPS/MFS according to the TCGA database.

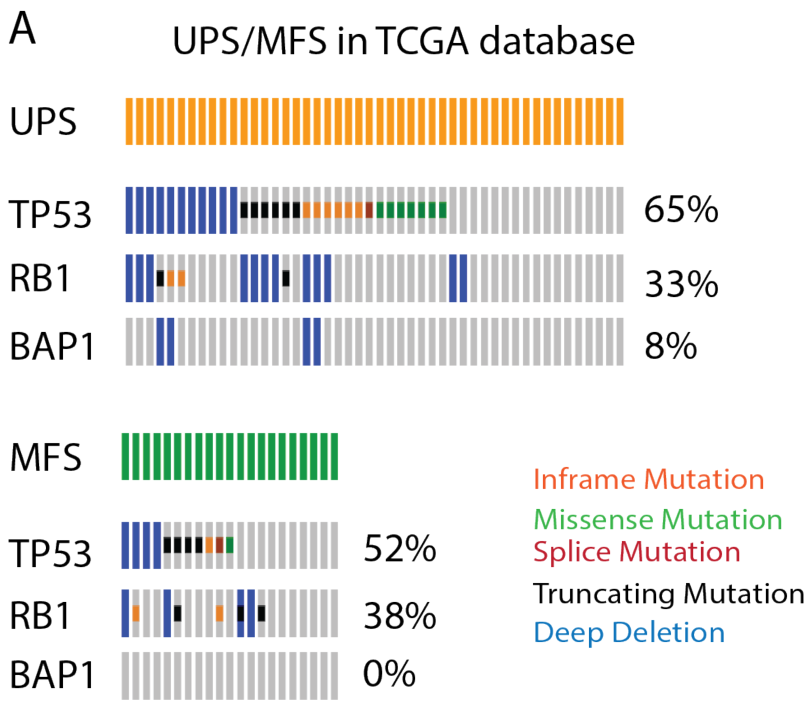

**Supplementary Figure 2**. Copy numbers of each sgRNA in the three libraries by sequencing.

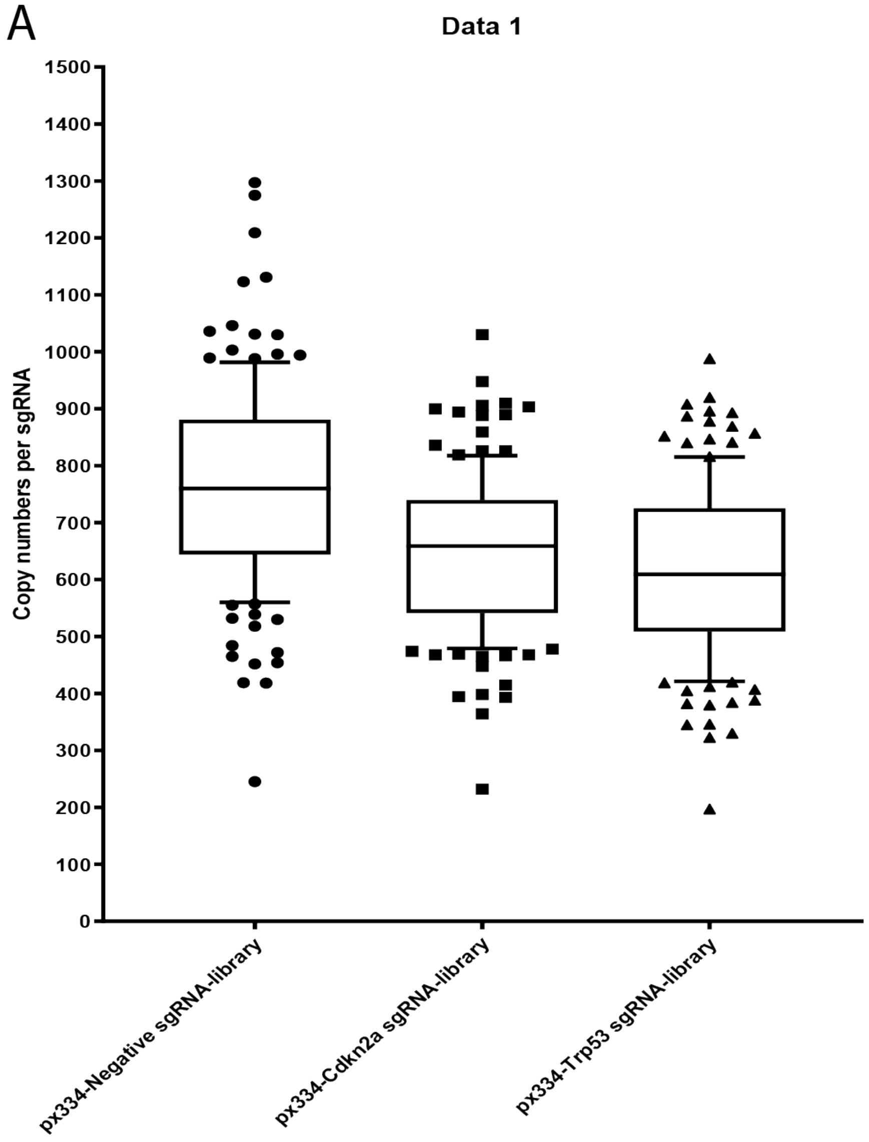

**Supplementary Figure 3**. Tumor free survival in Rosa26 $^{LoxP-Cas9/+}$ mice following IVE delivery of three different libraries. 5 mice per group.

A

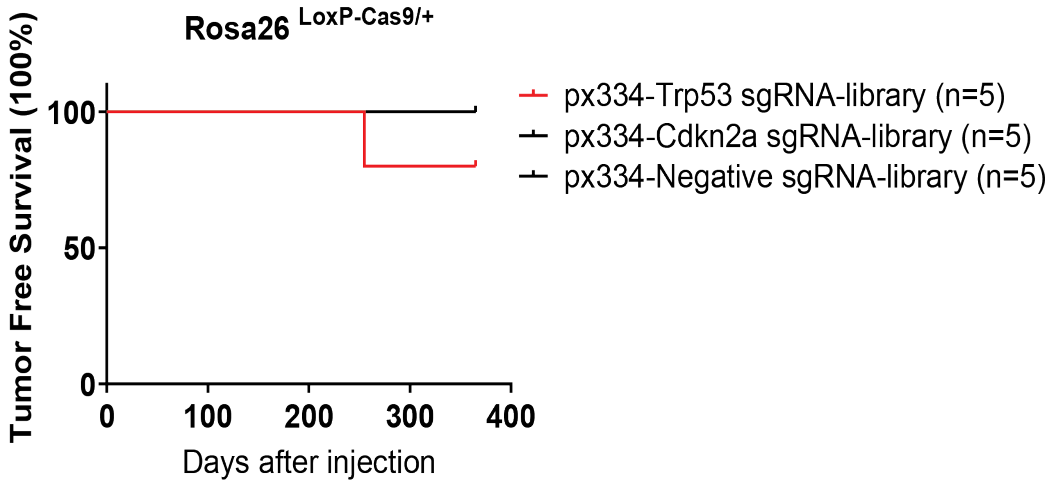

**Supplementary Figure 4**. (A) The Rosa26 $^{LoxP-Cas9/LoxP-Cas9}$ mice were intramuscularly injected (IVE) with 50 µg of mixed plasmids containing px333-Trp53-Pten sgRNAs and the px334-Trp53 sgRNA-library at a 1-1 ratio.

A

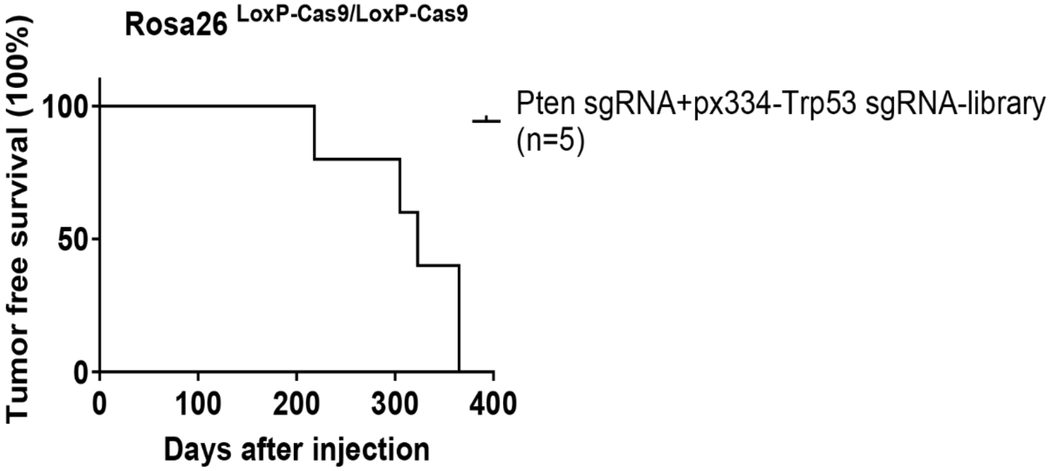

**Supplementary Figure 5**. (A) T7EI assay was performed to detect Indels in tumor DNAs induced by the px334-Trp53 sgRNA-library. (B) Sanger sequencing with Inference of CRISPR Edits (ICE) analysis was used to calculate the percentage of Indels in tumor S1 induced by the pX334-Trp53 sgRNA-library. (C) Next generation sequencing analysis was used to calculate the percentage of Indels in tumor S7, S11, and S13 induced by the pX334-Trp53 sgRNA-library.

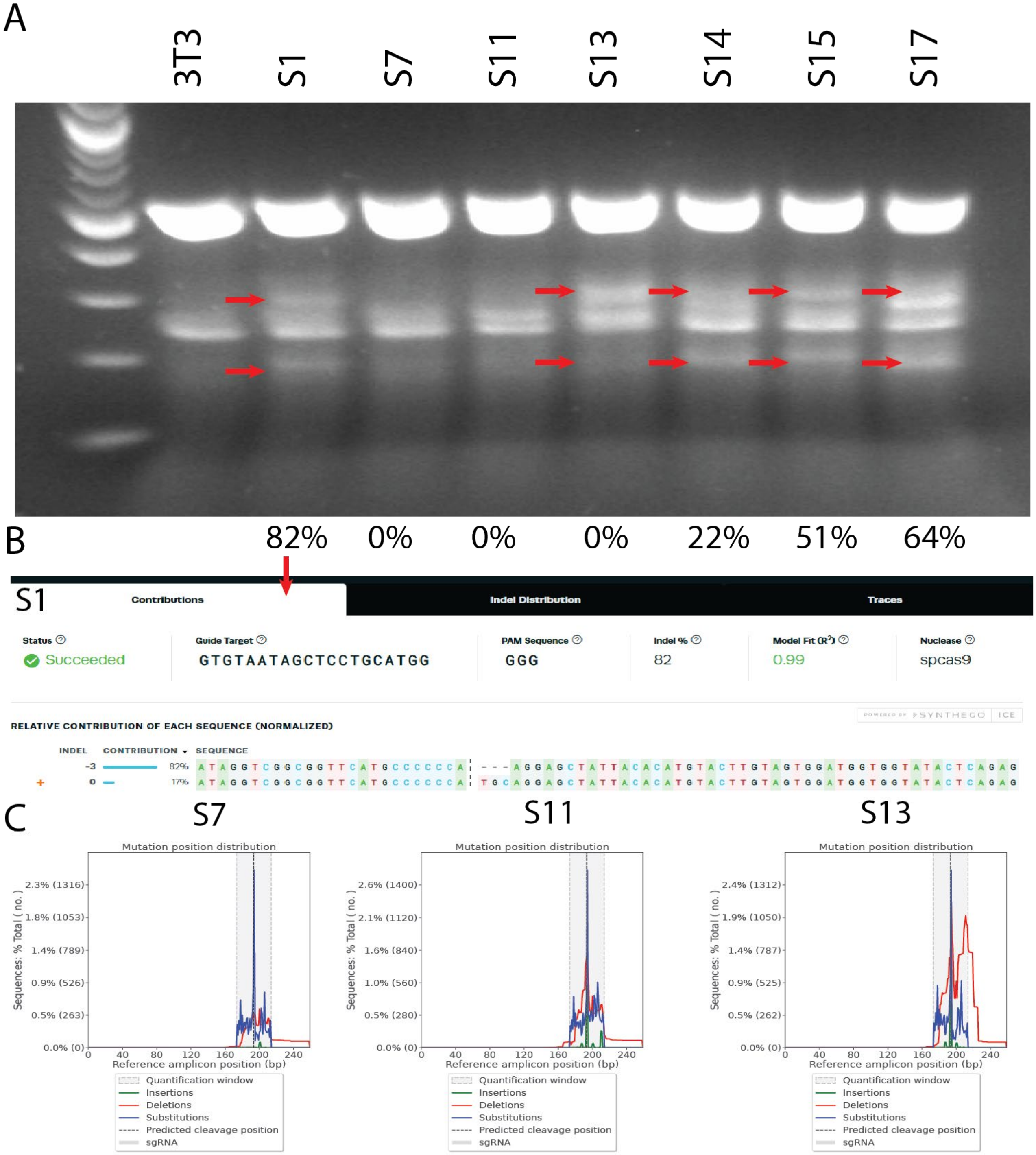

**Supplementary Figure 6**. No tumors were detected in mice IVE of plasmids containing either Trp53-Mllt3 sgRNAs, Trp53-Crebbp sgRNAs, Trp53-Pbrm1 sgRNAs, Trp53-Cysltr2, Trp53-Mst1r, or Trp53-Fat1 sgRNAs.

A

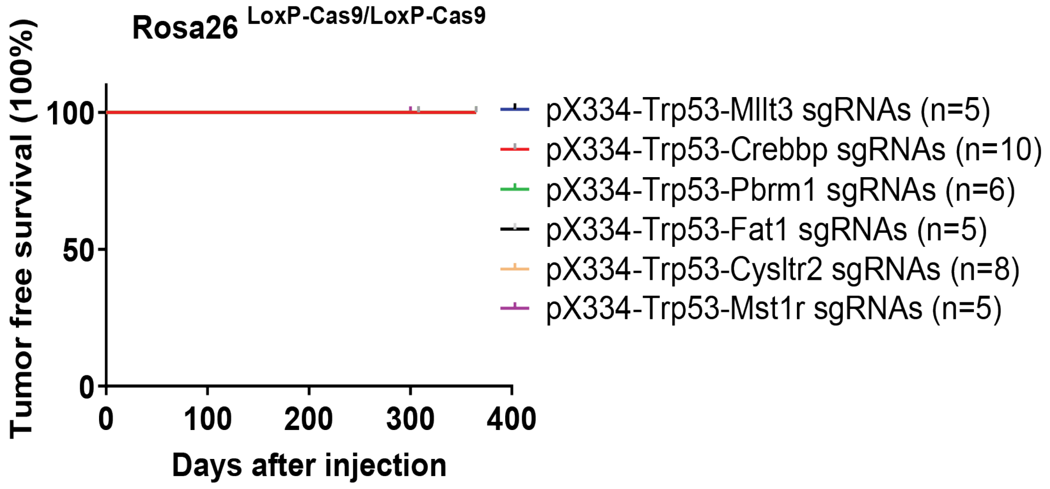

**Supplementary Figure 7**. H&E staining of mouse tumors driven by various gene mutations shows that these tumors are similar to each other. Scale bar = 50 μm.

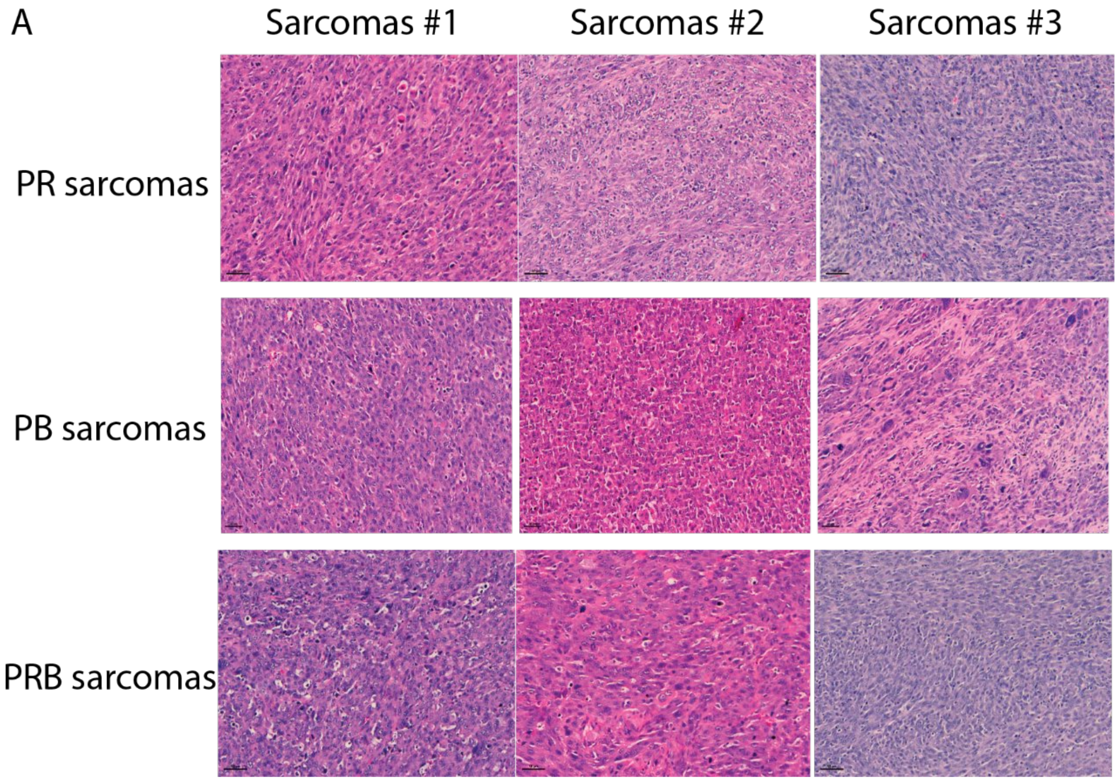

**Supplementary Figure 8**. Immunohistochemistry with CD68 in various mouse sarcoma tissue sections showed abundant $CD68^{+}$ macrophages in the tumor. Scale bar = 100 μm.

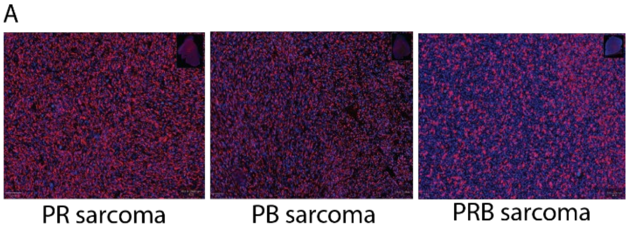

**Supplementary Figure 9**. (A) FC showed that there are no differences in the percentage of PD-1$^{+}$CD8$^{+}$ T cells between PR CRISPR large, PB CRISPR large and small tumors.

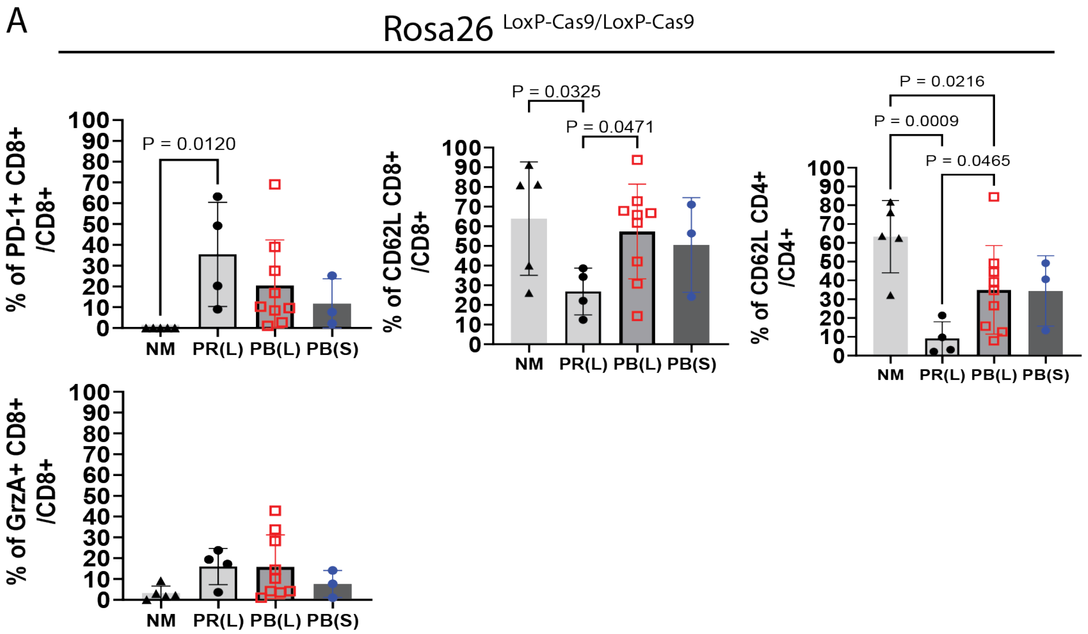

**Supplementary Figure 10**. (A) There are no differences in the percentage of CD11b$^{+}$ cells between PR CRISPR large, and PB CRISPR large tumors. However, the percentage of CD11b$^{+}$ cells is significantly lower in PB CRISPR early tumors compared to PB CRISPR large tumors. (B) Flow cytometry gating strategy to identify PMN-MDSCs, M-MDSCs, and TAMs. (C) There are no differences in the percentage of PMN-MDSCs, M-MDSCs, or TAMs between PB CRISPR large and small tumors. (D) There is a significantly low percentage of PD-L1$^{+}$ M-MDSCs in PRB Cre/CRISPR tumors compared to PR Cre tumors.

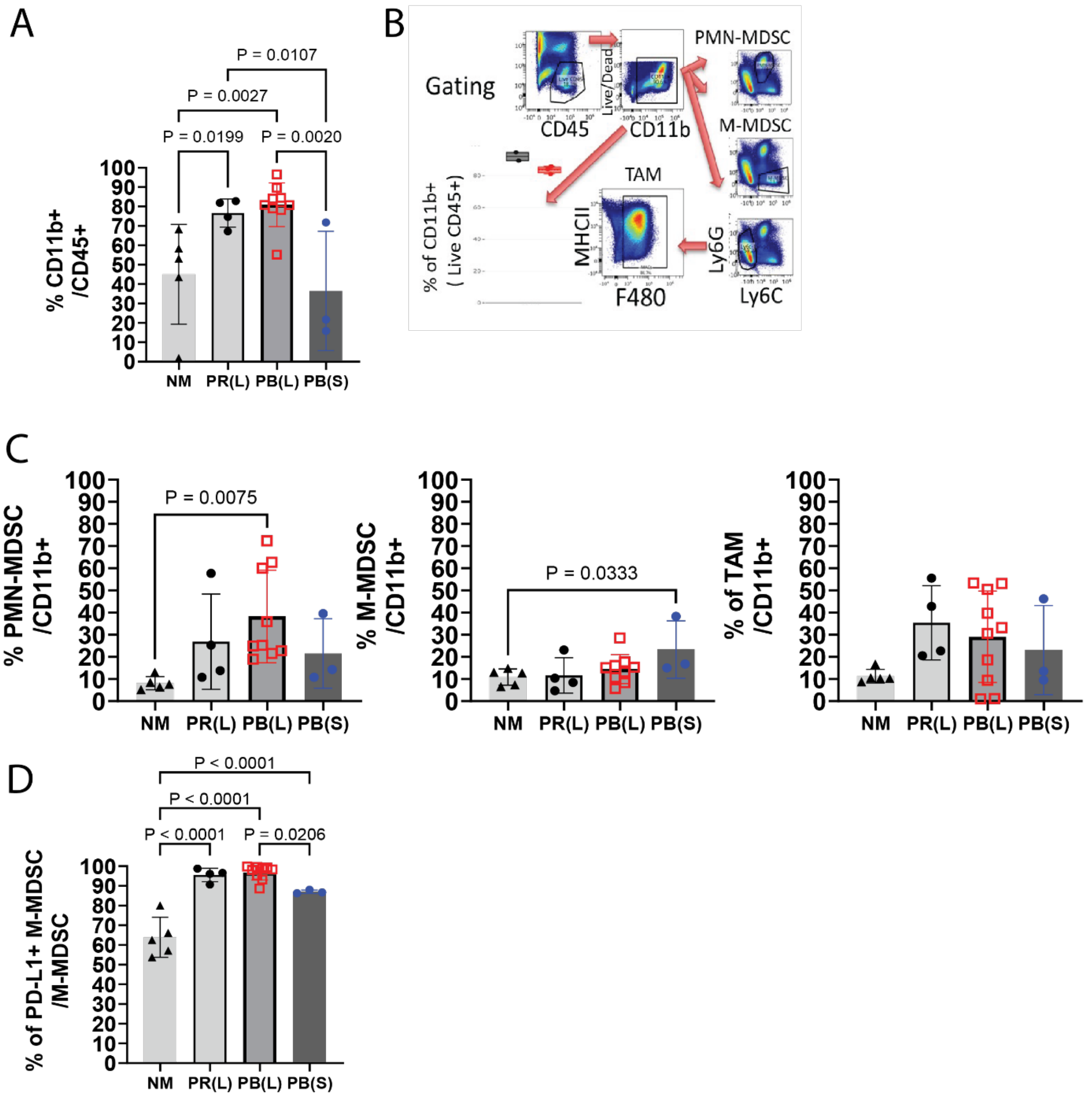

**Supplementary Figure 11**. (A) Western blot analysis to determine the knockout of Trp53 and Rb1 in PR cell lines. (B) Western blot analysis to determine the knockout of Trp53 and Bap1 in PB cell lines.

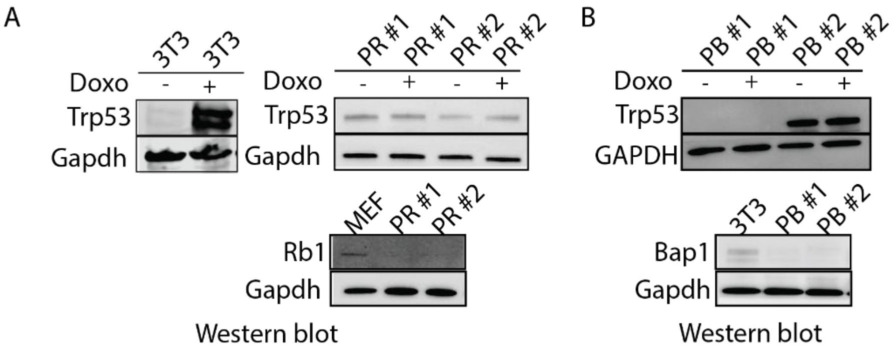

**Supplementary Figure 12**. (A) Both PR and PB mouse sarcoma cells are sensitive to two PARP inhibitors, niraparib and talazoparib, in a dose-dependent manner in vitro. (B) Niraparib significantly slows PR sarcoma growth in subcutaneous nude mice, while niraparib has no impact on PB sarcoma growth in subcutaneous nude mice. 5 mice in each group.

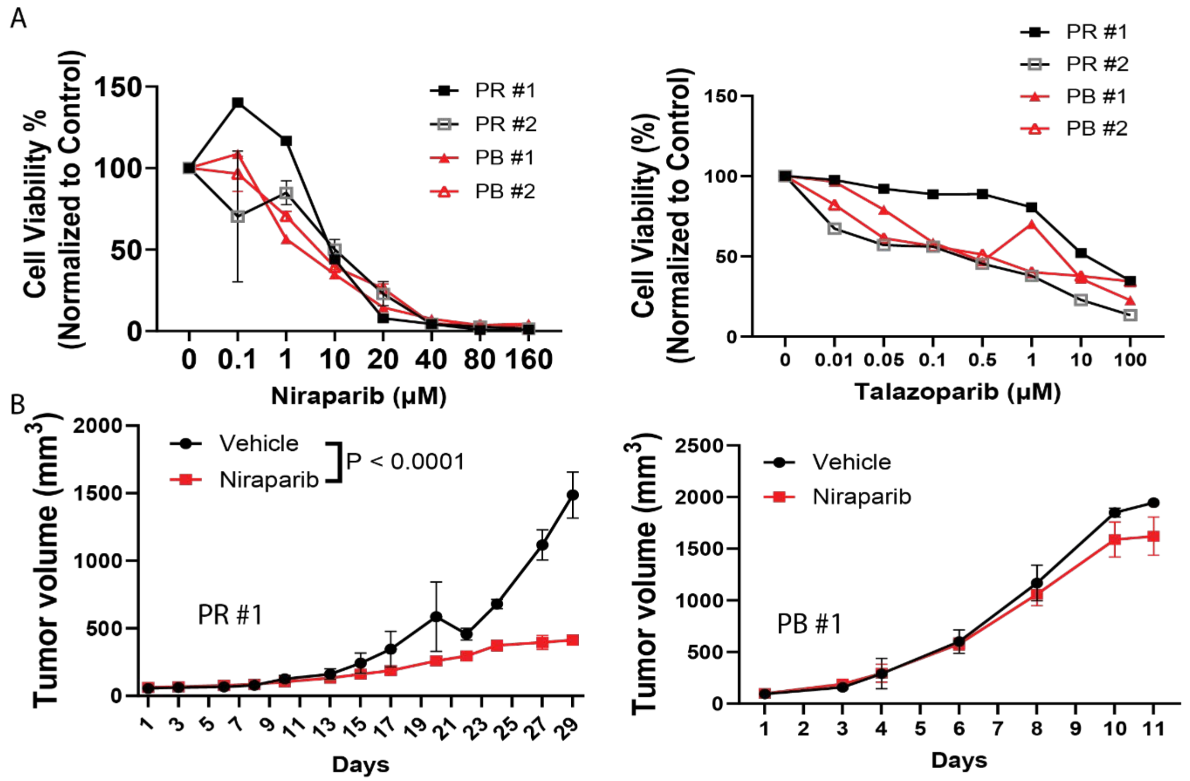

**Supplementary Figure 13**. There are no differences in the percentage of $CD11b^+$ cells between vehicle and volasertib treatment group.

A

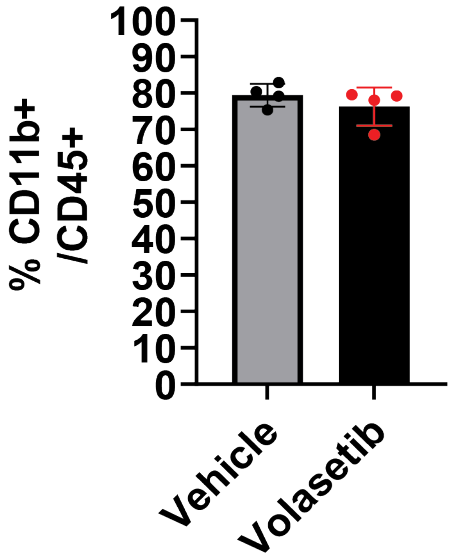

**Supplementary Figure 14**. Mutation profiling of PR-DUB complex genes in human UPS according to the TCGA database.

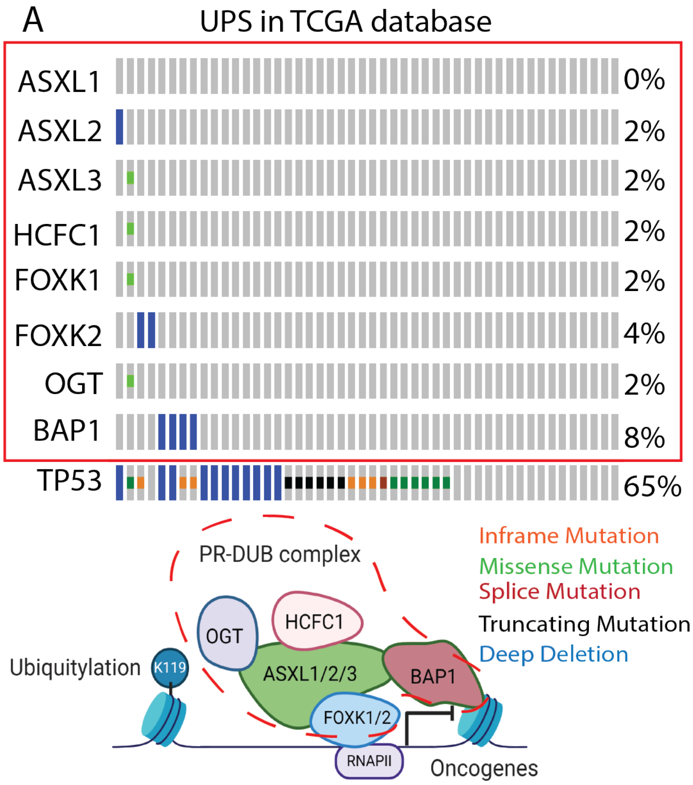

**Supplementary Figure 15**. Raw images of all western blotting are presented in this manuscript.

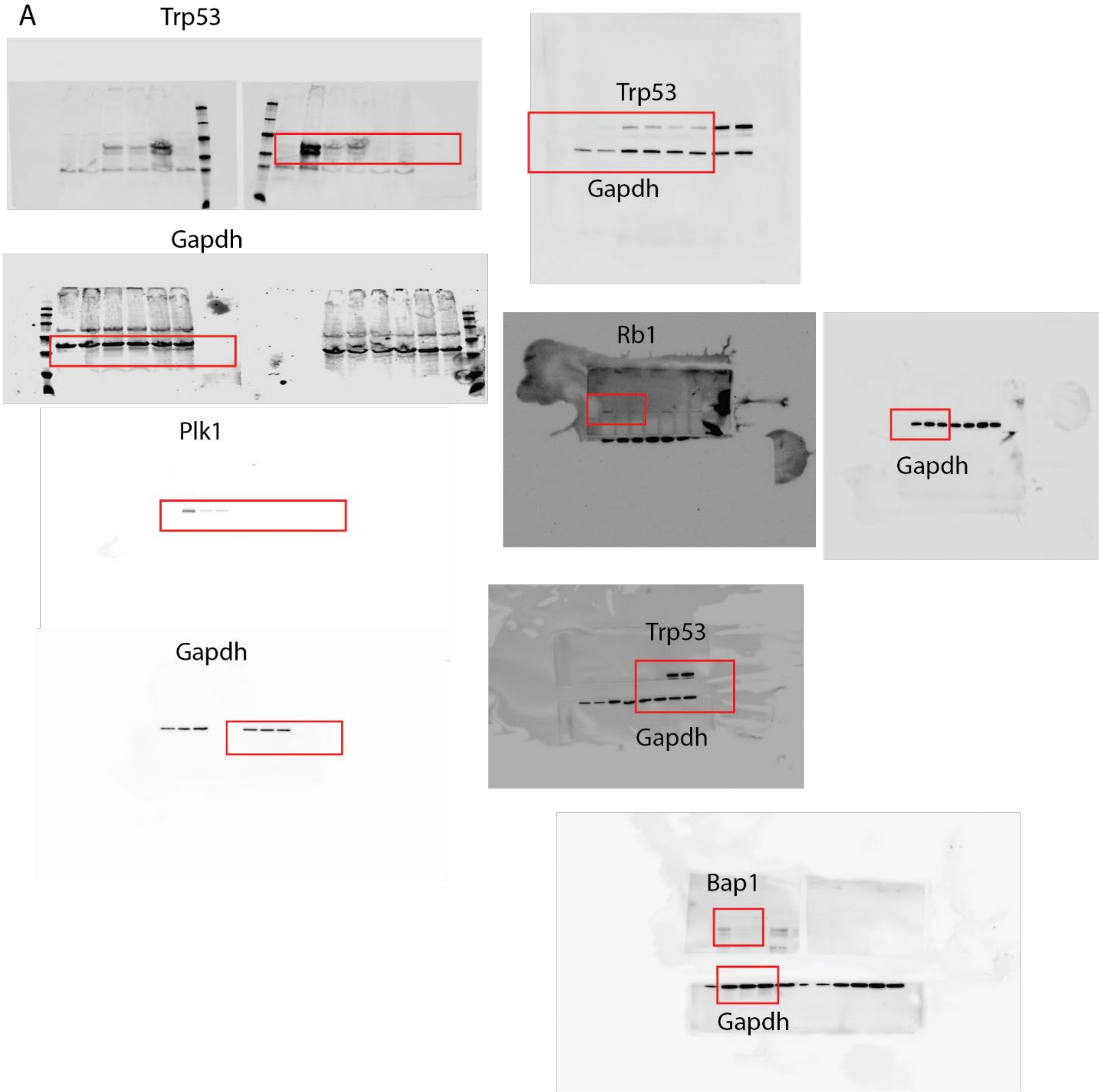

Supplementary Table 1. Top 35 genes plus TP53 and CDKN2A that are loss-of-function mutated or have copy number alterations in human UPSs and MFSs.

| Gene | | Copy Change | CN deletion TCGA | TCGA driver mut | CN deletion Genie | GENIE Frameshift mutations | GENIE other Driver mutations | Total # of samples with gene Alterations | Percentage of samples with Alterations | Total Number of UPS+MFS Samples |
|---|---|---|---|---|---|---|---|---|---|---|
| TP53 | 17p13.1 | DEL | 15 | 20 | 10 | 9 | 15 | 69 | 0.56097561 | 123 |
| RB1 | 13q14.2 | DEL | 16 | 5 | 12 | 6 | 1 | 40 | 0.32520325 | 123 |
| CYSLTR2 | 13q14.2 | DEL | 13 | | | | | 13 | 0.10569106 | 123 |
| CDKN2A | 9p21 | DEL | 14 | 1 | 10 | 3 | 0 | 28 | 0.22764228 | 123 |
| ATRX | Xq21.1 | DEL | 3 | 16 | 1 | 7 | 0 | 27 | 0.2195122 | 123 |
| D2HGDH | 2q37.3 | DEL | 10 | | | | | 10 | 0.08130081 | 123 |
| SNED1 | 2q37.3 | DEL | 10 | | | | | 10 | 0.08130081 | 123 |
| RHOA | 3p21.3 | DEL | 4 | 14 | 1 | 0 | 0 | 19 | 0.15447154 | 123 |
| CDKN2B | 9p21 | DEL | 14 | 0 | 9 | 0 | 0 | 23 | 0.18699187 | 123 |
| MTAP | 9p21 | DEL | 8 | | | | | 8 | 0.06504065 | 123 |
| HDAC4 | 2q37.3 | DEL | 8 | | | | | 8 | 0.06504065 | 123 |
| PASK | 2q37.3 | DEL | 7 | | | | | 7 | 0.05691057 | 123 |
| ACKR3 | 2q37.3 | DEL | 7 | | | | | 7 | 0.05691057 | 123 |
| ANO7 | | | 7 | | | | | 7 | 0.04605263 | 152 |
| FAT1 | 4q35 | DEL | 6 | 3 | 1 | 1 | 0 | 11 | 0.08943089 | 123 |
| PDCD1 | 2q37.3 | DEL | 11 | | | | | 11 | 0.08943089 | 123 |
| WWOX | 16q23 | DEL | 5 | | | | | 5 | 0.04065041 | 123 |
| DNAH12 | 3p14.3 | DEL | 4 | | | | | 4 | 0.03252033 | 123 |
| FHIT | 3p14.2 | DEL | 4 | | | | | 4 | 0.03252033 | 123 |
| INPP5D | 2q37.1 | DEL | 4 | | | | | 4 | 0.03252033 | 123 |
| MLLT3 | 9p22 | DEL | 4 | | | | | 4 | 0.03252033 | 123 |
| RYR1 | | | 0 | 4 | 0 | 0 | 0 | 4 | 0.03252033 | 123 |
| NUMBL | | | 1 | 3 | 0 | 0 | 0 | 4 | 0.03252033 | 123 |
| PARP3 | 3p21.2 | DEL | 4 | | | | | 4 | 0.03252033 | 123 |
| MST1R | 3p21.3 | DEL | 4 | | 2 | 0 | 0 | 6 | 0.04878049 | 123 |
| CSDMD1 | | | 4 | | | | | 4 | 0.02631579 | 152 |
| MST1 | 3p21 | DEL | 4 | | 1 | 0 | 0 | 5 | 0.04065041 | 123 |
| KDM4C | 9p24.1 | DEL | 3 | | | | | 3 | 0.02439024 | 123 |
| PSIP1 | 9p22.3 | DEL | 3 | | | | | 3 | 0.02439024 | 123 |
| MYT1L | | | 2 | 1 | | | | 3 | 0.01973684 | 152 |
| PBRM1 | 3p21 | DEL | 4 | | 1 | 0 | 0 | 5 | 0.04065041 | 123 |
| BAP1 | 3p21.1 | DEL | 4 | 0 | 1 | 0 | 0 | 5 | 0.04065041 | 123 |
| PDCD1LG2 | 9p24.2 | DEL | 3 | | | | | 3 | 0.02439024 | 123 |
| JAK2 | 9p24 | DEL | 3 | | 0 | 0 | 1 | 4 | 0.03252033 | 123 |
| IRS1 | 2q36 | DEL | 3 | | | | | 3 | 0.02439024 | 123 |
| CD274 | 9p24 | DEL | 3 | | | | | 3 | 0.02439024 | 123 |
| CREBBP | | | 0 | 2 | 0 | 0 | 0 | 2 | 0.01626016 | 123 |

Supplementary Table 2. Next generation sequencing detected InDels in the Trp53 locus targeted by CRISPR/Cas9.

| SAMPLE | READS.ALIGNED | Unmodified Reads | Unmodified (%) | Mutant Reads | InDel (%) |
|---|---|---|---|---|---|
| S7 | 57699 | 52529 | 91.04 | 5170 | 8.96 |
| S11 | 53615 | 47621 | 88.82 | 5994 | 11.18 |
| S13 | 55304 | 48583 | 87.85 | 6721 | 12.15 |
| S14 | 65588 | 29444 | 44.89 | 36144 | 55.11 |
| S15 | 64401 | 27771 | 43.12 | 36630 | 56.88 |
| S17 | 94934 | 30627 | 32.26 | 64307 | 67.74 |

Supplementary Table 3. The rate of metastasis detected in the mice bearing different gene mutation driven tumors.

| Genotype | | Tumors | Mets (Lung, otherwise specified) | Lung Met Rate | Note |
|---|---|---|---|---|---|
| Trp53/Rb1 CRISPR | | 11 | 1 | 9.09 | |
| Trp53/Rb1 Cre | | 2 | 1 | 50.00 | |
| Trp53/Bap1 CRISPR | | 16 | 4 | 25.00 | |
| Trp53/Rb1/Bap1 Cre/ | | 6 | 1 | 16.67 | |

Supplementary Table 4. Oligonucleotides.

| Number | Target Gene ID | Target Gene Symbol | sgRNA Target Sequence | FWD oligoes | REV oligoes |
|---|---|---|---|---|---|
| 73664 | 404545 | Ano7 | ACTGGCAGGAGTAATACTCG | **CACCG**ACTGGCAGGAGTAATACTCG | **AAAC**CGAGTATTACTCCTGCCAGT**C** |
| 73665 | 404545 | Ano7 | CCATGTGCGAAGGTACTTCG | **CACCG**CCATGTGCGAAGGTACTTCG | **AAAC**CGAAGTACCTTCGCACATGG**C** |
| 73666 | 404545 | Ano7 | GAAGGCGATATACACGGGCG | **CACCG**GAAGGCGATATACACGGGCG | **AAAC**CGCCCGTGTATATCGCCTTC**C** |
| 73667 | 404545 | Ano7 | ATAGATGGCTTACCAATTGG | **CACCG**ATAGATGGCTTACCAATTGG | **AAAC**CCAATTGGTAAGCCATCTAT**C** |
| 46056 | 94109 | Csmd1 | GTACTTACCAATGCACAAAG | **CACCG**GTACTTACCAATGCACAAAG | **AAAC**CTTTGTGCATTGGTAAGTAC**C** |
| 46057 | 94109 | Csmd1 | GTCACACACTCAATGGACGA | **CACCG**GTCACACACTCAATGGACGA | **AAAC**TCGTCCATTGAGTGTGTGAC**C** |
| 46058 | 94109 | Csmd1 | CGAACTGAAACCCAATTCTG | **CACCG**CGAACTGAAACCCAATTCTG | **AAAC**CAGAATTGGGTTTCAGTTCG**C** |
| 46059 | 94109 | Csmd1 | GTGGTACTCTCCTATGAGTG | **CACCG**GTGGTACTCTCCTATGAGTG | **AAAC**CACTCATAGGAGAGTACCAC**C** |
| 11314 | 17933 | Myt1l | TCCCCCAAAGGATATGACGA | **CACCG**TCCCCCAAAGGATATGACGA | **AAAC**TCGTCATATCCTTTGGGGGA**C** |
| 11315 | 17933 | Myt1l | CTCATAACATTCGTCTACTG | **CACCG**CTCATAACATTCGTCTACTG | **AAAC**CAGTAGACGAATGTTATGAG**C** |
| 11316 | 17933 | Myt1l | GTGACAGAAGTTATGCTGAG | **CACCG**GTGACAGAAGTTATGCTGAG | **AAAC**CTCAGCATAACTTCTGTCAC**C** |
| 11317 | 17933 | Myt1l | GCGGGTAAAGCCCAGTTACG | **CACCG**GCGGGTAAAGCCCAGTTACG | **AAAC**CGTAACTGGGCTTTACCCGC**C** |

### Supplementary Table 5. Oligonucleotide pools for synthesis.

| Number | Target Ge | Target Gene Symb | Left arm sequence | sgRNA Target Sequence | Right arm sequence | Oligonucleotide pools for synthesis |
|---|---|---|---|---|---|---|
| 14482 | 19645 | Rb1 | ATATATCTTGTGGAAAGGACGAAACACCG | AAATGATACGAGGATTATCG | GTTTTAGAGCTAGAAATAGCAAGTTAAAA | ATATATCTTGTGGAAAGGACGAAACACCGAAATGATACGAGGATTATCGGTTTTAGAGCTAGAAATAGCAAGTTAAAA |
| 14483 | 19645 | Rb1 | ATATATCTTGTGGAAAGGACGAAACACCG | AGAGAAGTTTGCTAACGCTG | GTTTTAGAGCTAGAAATAGCAAGTTAAAA | ATATATCTTGTGGAAAGGACGAAACACCGAGAGAAGTTTGCTAACGCTGGTTTTAGAGCTAGAAATAGCAAGTTAAAA |
| 14484 | 19645 | Rb1 | ATATATCTTGTGGAAAGGACGAAACACCG | TAAGTACGTTCAGAATCCAC | GTTTTAGAGCTAGAAATAGCAAGTTAAAA | ATATATCTTGTGGAAAGGACGAAACACCGTAAGTACGTTCAGAATCCACGTTTTAGAGCTAGAAATAGCAAGTTAAAA |
| 14485 | 19645 | Rb1 | ATATATCTTGTGGAAAGGACGAAACACCG | GCAGTATGGTTACCCTGGAG | GTTTTAGAGCTAGAAATAGCAAGTTAAAA | ATATATCTTGTGGAAAGGACGAAACACCGGCAGTATGGTTACCCTGGAGGTTTTAGAGCTAGAAATAGCAAGTTAAAA |
| 35366 | 70086 | Cysltr2 | ATATATCTTGTGGAAAGGACGAAACACCG | AAACCCATATGATCCCACAG | GTTTTAGAGCTAGAAATAGCAAGTTAAAA | ATATATCTTGTGGAAAGGACGAAACACCGAAACCCATATGATCCCACAGGTTTTAGAGCTAGAAATAGCAAGTTAAAA |
| 35367 | 70086 | Cysltr2 | ATATATCTTGTGGAAAGGACGAAACACCG | AACCACATTGCAGTAGCGGT | GTTTTAGAGCTAGAAATAGCAAGTTAAAA | ATATATCTTGTGGAAAGGACGAAACACCGAACCACATTGCAGTAGCGGTGTTTTAGAGCTAGAAATAGCAAGTTAAAA |
| 35368 | 70086 | Cysltr2 | ATATATCTTGTGGAAAGGACGAAACACCG | TATGGAAAAGCCATTTCCCA | GTTTTAGAGCTAGAAATAGCAAGTTAAAA | ATATATCTTGTGGAAAGGACGAAACACCGTATGGAAAAGCCATTTCCCAGTTTTAGAGCTAGAAATAGCAAGTTAAAA |
| 35369 | 70086 | Cysltr2 | ATATATCTTGTGGAAAGGACGAAACACCG | GATGAATAGAAAATCGGAAG | GTTTTAGAGCTAGAAATAGCAAGTTAAAA | ATATATCTTGTGGAAAGGACGAAACACCGGATGAATAGAAAATCGGAAGGTTTTAGAGCTAGAAATAGCAAGTTAAAA |
| 1049 | 11848 | Rhoa | ATATATCTTGTGGAAAGGACGAAACACCG | AAAACACATCAATATAACAT | GTTTTAGAGCTAGAAATAGCAAGTTAAAA | ATATATCTTGTGGAAAGGACGAAACACCGAAAACACATCAATATAACATGTTTTAGAGCTAGAAATAGCAAGTTAAAA |
| 1050 | 11848 | Rhoa | ATATATCTTGTGGAAAGGACGAAACACCG | CTATGTGGCGGATATCGAGG | GTTTTAGAGCTAGAAATAGCAAGTTAAAA | ATATATCTTGTGGAAAGGACGAAACACCGCTATGTGGCGGATATCGAGGGTTTTAGAGCTAGAAATAGCAAGTTAAAA |
| 1051 | 11848 | Rhoa | ATATATCTTGTGGAAAGGACGAAACACCG | GCTACTCACCTAAACTATCA | GTTTTAGAGCTAGAAATAGCAAGTTAAAA | ATATATCTTGTGGAAAGGACGAAACACCGGCTACTCACCTAAACTATCAGTTTTAGAGCTAGAAATAGCAAGTTAAAA |
| 1052 | 11848 | Rhoa | ATATATCTTGTGGAAAGGACGAAACACCG | CAGCAAGGACCAGTTCCCAG | GTTTTAGAGCTAGAAATAGCAAGTTAAAA | ATATATCTTGTGGAAAGGACGAAACACCGCAGCAAGGACCAGTTCCCAGGTTTTAGAGCTAGAAATAGCAAGTTAAAA |
| 46424 | 98314 | D2hgdh | ATATATCTTGTGGAAAGGACGAAACACCG | GGCTCACAGAGCATACTCCG | GTTTTAGAGCTAGAAATAGCAAGTTAAAA | ATATATCTTGTGGAAAGGACGAAACACCGGGCTCACAGAGCATACTCCGGTTTTAGAGCTAGAAATAGCAAGTTAAAA |
| 46425 | 98314 | D2hgdh | ATATATCTTGTGGAAAGGACGAAACACCG | AGGCCTGGGTGGGCACACAA | GTTTTAGAGCTAGAAATAGCAAGTTAAAA | ATATATCTTGTGGAAAGGACGAAACACCGAGGCCTGGGTGGGCACACAAGTTTTAGAGCTAGAAATAGCAAGTTAAAA |
| 46426 | 98314 | D2hgdh | ATATATCTTGTGGAAAGGACGAAACACCG | GGATACTGAATGCATGCAGT | GTTTTAGAGCTAGAAATAGCAAGTTAAAA | ATATATCTTGTGGAAAGGACGAAACACCGGGATACTGAATGCATGCAGTGTTTTAGAGCTAGAAATAGCAAGTTAAAA |
| 46427 | 98314 | D2hgdh | ATATATCTTGTGGAAAGGACGAAACACCG | GACTGGCTGAAGACCGTCCG | GTTTTAGAGCTAGAAATAGCAAGTTAAAA | ATATATCTTGTGGAAAGGACGAAACACCGGACTGGCTGAAGACCGTCCGGTTTTAGAGCTAGAAATAGCAAGTTAAAA |
| 29438 | 66902 | Mtap | ATATATCTTGTGGAAAGGACGAAACACCG | TGTGGATACTCCATTCGGCA | GTTTTAGAGCTAGAAATAGCAAGTTAAAA | ATATATCTTGTGGAAAGGACGAAACACCGTGTGGATACTCCATTCGGCAGTTTTAGAGCTAGAAATAGCAAGTTAAAA |
| 29439 | 66902 | Mtap | ATATATCTTGTGGAAAGGACGAAACACCG | GGACAATAGTCACAATTGAG | GTTTTAGAGCTAGAAATAGCAAGTTAAAA | ATATATCTTGTGGAAAGGACGAAACACCGGGACAATAGTCACAATTGAGGTTTTAGAGCTAGAAATAGCAAGTTAAAA |
| 29440 | 66902 | Mtap | ATATATCTTGTGGAAAGGACGAAACACCG | GGCAAAACGGTTCAGCCATG | GTTTTAGAGCTAGAAATAGCAAGTTAAAA | ATATATCTTGTGGAAAGGACGAAACACCGGGCAAAACGGTTCAGCCATGGTTTTAGAGCTAGAAATAGCAAGTTAAAA |
| 29441 | 66902 | Mtap | ATATATCTTGTGGAAAGGACGAAACACCG | GCCTTCAAAAGTCAACTACC | GTTTTAGAGCTAGAAATAGCAAGTTAAAA | ATATATCTTGTGGAAAGGACGAAACACCGGCCTTCAAAAGTCAACTACCGTTTTAGAGCTAGAAATAGCAAGTTAAAA |
| 2669 | 12579 | Cdkn2b | ATATATCTTGTGGAAAGGACGAAACACCG | GTGCACGACGCAGCTCGGGA | GTTTTAGAGCTAGAAATAGCAAGTTAAAA | ATATATCTTGTGGAAAGGACGAAACACCGGTGCACGACGCAGCTCGGGAGTTTTAGAGCTAGAAATAGCAAGTTAAAA |
| 2670 | 12579 | Cdkn2b | ATATATCTTGTGGAAAGGACGAAACACCG | GTCGTGCACAGGTCTGGTAA | GTTTTAGAGCTAGAAATAGCAAGTTAAAA | ATATATCTTGTGGAAAGGACGAAACACCGGTCGTGCACAGGTCTGGTAAGTTTTAGAGCTAGAAATAGCAAGTTAAAA |
| 2671 | 12579 | Cdkn2b | ATATATCTTGTGGAAAGGACGAAACACCG | CAGGGCGTTGGGATCTGCGC | GTTTTAGAGCTAGAAATAGCAAGTTAAAA | ATATATCTTGTGGAAAGGACGAAACACCGCAGGGCGTTGGGATCTGCGCGTTTTAGAGCTAGAAATAGCAAGTTAAAA |
| 2672 | 12579 | Cdkn2b | ATATATCTTGTGGAAAGGACGAAACACCG | GGATTGGGCGCCTCCCGAAG | GTTTTAGAGCTAGAAATAGCAAGTTAAAA | ATATATCTTGTGGAAAGGACGAAACACCGGGATTGGGCGCCTCCCGAAGGTTTTAGAGCTAGAAATAGCAAGTTAAAA |
| 68212 | 269224 | Pask | ATATATCTTGTGGAAAGGACGAAACACCG | GTACCGAACTATCATATGTG | GTTTTAGAGCTAGAAATAGCAAGTTAAAA | ATATATCTTGTGGAAAGGACGAAACACCGGTACCGAACTATCATATGTGGTTTTAGAGCTAGAAATAGCAAGTTAAAA |
| 68213 | 269224 | Pask | ATATATCTTGTGGAAAGGACGAAACACCG | CTGTCCTTAACTGAAACTAG | GTTTTAGAGCTAGAAATAGCAAGTTAAAA | ATATATCTTGTGGAAAGGACGAAACACCGCTGTCCTTAACTGAAACTAGGTTTTAGAGCTAGAAATAGCAAGTTAAAA |
| 68214 | 269224 | Pask | ATATATCTTGTGGAAAGGACGAAACACCG | TACCACATCTGAACTCAGCG | GTTTTAGAGCTAGAAATAGCAAGTTAAAA | ATATATCTTGTGGAAAGGACGAAACACCGTACCACATCTGAACTCAGCGGTTTTAGAGCTAGAAATAGCAAGTTAAAA |
| 68215 | 269224 | Pask | ATATATCTTGTGGAAAGGACGAAACACCG | TGCAGTGGTGTTCGGCACAG | GTTTTAGAGCTAGAAATAGCAAGTTAAAA | ATATATCTTGTGGAAAGGACGAAACACCGTGCAGTGGTGTTCGGCACAGGTTTTAGAGCTAGAAATAGCAAGTTAAAA |
| 52568 | 208777 | Sned1 | ATATATCTTGTGGAAAGGACGAAACACCG | CATCGATTCTAAACGCCCAG | GTTTTAGAGCTAGAAATAGCAAGTTAAAA | ATATATCTTGTGGAAAGGACGAAACACCGCATCGATTCTAAACGCCCAGGTTTTAGAGCTAGAAATAGCAAGTTAAAA |
| 52569 | 208777 | Sned1 | ATATATCTTGTGGAAAGGACGAAACACCG | GGTGGACTGTGGTCACCCTG | GTTTTAGAGCTAGAAATAGCAAGTTAAAA | ATATATCTTGTGGAAAGGACGAAACACCGGGTGGACTGTGGTCACCCTGGTTTTAGAGCTAGAAATAGCAAGTTAAAA |
| 52570 | 208777 | Sned1 | ATATATCTTGTGGAAAGGACGAAACACCG | AGGTGATGTCTACTACCGGG | GTTTTAGAGCTAGAAATAGCAAGTTAAAA | ATATATCTTGTGGAAAGGACGAAACACCGAGGTGATGTCTACTACCGGGGTTTTAGAGCTAGAAATAGCAAGTTAAAA |
| 52571 | 208777 | Sned1 | ATATATCTTGTGGAAAGGACGAAACACCG | CGCACGTGTAGGAGTCCTCG | GTTTTAGAGCTAGAAATAGCAAGTTAAAA | ATATATCTTGTGGAAAGGACGAAACACCGCGCACGTGTAGGAGTCCTCGGTTTTAGAGCTAGAAATAGCAAGTTAAAA |
| 5389 | 14107 | Fat1 | ATATATCTTGTGGAAAGGACGAAACACCG | GTGCGCGCCCATCATAACCG | GTTTTAGAGCTAGAAATAGCAAGTTAAAA | ATATATCTTGTGGAAAGGACGAAACACCGGTGCGCGCCCATCATAACCGGTTTTAGAGCTAGAAATAGCAAGTTAAAA |
| 5390 | 14107 | Fat1 | ATATATCTTGTGGAAAGGACGAAACACCG | ACGACCTCGGTATACCCCAG | GTTTTAGAGCTAGAAATAGCAAGTTAAAA | ATATATCTTGTGGAAAGGACGAAACACCGACGACCTCGGTATACCCCAGGTTTTAGAGCTAGAAATAGCAAGTTAAAA |
| 5391 | 14107 | Fat1 | ATATATCTTGTGGAAAGGACGAAACACCG | ACGGTCAAACGGTATCCCGG | GTTTTAGAGCTAGAAATAGCAAGTTAAAA | ATATATCTTGTGGAAAGGACGAAACACCGACGGTCAAACGGTATCCCGGGTTTTAGAGCTAGAAATAGCAAGTTAAAA |
| 5392 | 14107 | Fat1 | ATATATCTTGTGGAAAGGACGAAACACCG | TCCGTCGCTCGAATCACCAG | GTTTTAGAGCTAGAAATAGCAAGTTAAAA | ATATATCTTGTGGAAAGGACGAAACACCGTCCGTCGCTCGAATCACCAGGTTTTAGAGCTAGAAATAGCAAGTTAAAA |
| 52552 | 208727 | Hdac4 | ATATATCTTGTGGAAAGGACGAAACACCG | TGTGGCTGCCAAATGACACA | GTTTTAGAGCTAGAAATAGCAAGTTAAAA | ATATATCTTGTGGAAAGGACGAAACACCGTGTGGCTGCCAAATGACACAGTTTTAGAGCTAGAAATAGCAAGTTAAAA |
| 52553 | 208727 | Hdac4 | ATATATCTTGTGGAAAGGACGAAACACCG | CCTACCTAGATCGGCTACCT | GTTTTAGAGCTAGAAATAGCAAGTTAAAA | ATATATCTTGTGGAAAGGACGAAACACCGCCTACCTAGATCGGCTACCTGTTTTAGAGCTAGAAATAGCAAGTTAAAA |
| 52554 | 208727 | Hdac4 | ATATATCTTGTGGAAAGGACGAAACACCG | CTTCTGTCTACAGACGAGCT | GTTTTAGAGCTAGAAATAGCAAGTTAAAA | ATATATCTTGTGGAAAGGACGAAACACCGCTTCTGTCTACAGACGAGCTGTTTTAGAGCTAGAAATAGCAAGTTAAAA |
| 52555 | 208727 | Hdac4 | ATATATCTTGTGGAAAGGACGAAACACCG | TCACTTACCCATACCAGTAG | GTTTTAGAGCTAGAAATAGCAAGTTAAAA | ATATATCTTGTGGAAAGGACGAAACACCGTCACTTACCCATACCAGTAGGTTTTAGAGCTAGAAATAGCAAGTTAAAA |
| 44764 | 80707 | Wwox | ATATATCTTGTGGAAAGGACGAAACACCG | CTTCGTCGGATTATCGTCCA | GTTTTAGAGCTAGAAATAGCAAGTTAAAA | ATATATCTTGTGGAAAGGACGAAACACCGCTTCGTCGGATTATCGTCCAGTTTTAGAGCTAGAAATAGCAAGTTAAAA |
| 44765 | 80707 | Wwox | ATATATCTTGTGGAAAGGACGAAACACCG | CTTGGCAGTCTCGAACCCTG | GTTTTAGAGCTAGAAATAGCAAGTTAAAA | ATATATCTTGTGGAAAGGACGAAACACCGCTTGGCAGTCTCGAACCCTGGTTTTAGAGCTAGAAATAGCAAGTTAAAA |
| 44766 | 80707 | Wwox | ATATATCTTGTGGAAAGGACGAAACACCG | AAGAGGAAACGGGTCGCAGG | GTTTTAGAGCTAGAAATAGCAAGTTAAAA | ATATATCTTGTGGAAAGGACGAAACACCGAAGAGGAAACGGGTCGCAGGGTTTTAGAGCTAGAAATAGCAAGTTAAAA |
| 44767 | 80707 | Wwox | ATATATCTTGTGGAAAGGACGAAACACCG | ATTGCAGCACAAAGCCAAGG | GTTTTAGAGCTAGAAATAGCAAGTTAAAA | ATATATCTTGTGGAAAGGACGAAACACCGATTGCAGCACAAAGCCAAGGGTTTTAGAGCTAGAAATAGCAAGTTAAAA |
| 18966 | 22589 | Atrx | ATATATCTTGTGGAAAGGACGAAACACCG | GAATGGCCGTAAAAGTTCTG | GTTTTAGAGCTAGAAATAGCAAGTTAAAA | ATATATCTTGTGGAAAGGACGAAACACCGGAATGGCCGTAAAAGTTCTGGTTTTAGAGCTAGAAATAGCAAGTTAAAA |
| 18967 | 22589 | Atrx | ATATATCTTGTGGAAAGGACGAAACACCG | GACAATCTCAAAAAACGCGG | GTTTTAGAGCTAGAAATAGCAAGTTAAAA | ATATATCTTGTGGAAAGGACGAAACACCGGACAATCTCAAAAAACGCGGGTTTTAGAGCTAGAAATAGCAAGTTAAAA |
| 18968 | 22589 | Atrx | ATATATCTTGTGGAAAGGACGAAACACCG | GGGAGTTGCTCCATAGCAGG | GTTTTAGAGCTAGAAATAGCAAGTTAAAA | ATATATCTTGTGGAAAGGACGAAACACCGGGGAGTTGCTCCATAGCAGGGTTTTAGAGCTAGAAATAGCAAGTTAAAA |
| 18969 | 22589 | Atrx | ATATATCTTGTGGAAAGGACGAAACACCG | ACATACCTAACTGCTCTCTG | GTTTTAGAGCTAGAAATAGCAAGTTAAAA | ATATATCTTGTGGAAAGGACGAAACACCGACATACCTAACTGCTCTCTGGTTTTAGAGCTAGAAATAGCAAGTTAAAA |
| 12666 | 18566 | Pdcd1 | ATATATCTTGTGGAAAGGACGAAACACCG | CAATACAGGGATACCCACTA | GTTTTAGAGCTAGAAATAGCAAGTTAAAA | ATATATCTTGTGGAAAGGACGAAACACCGCAATACAGGGATACCCACTAGTTTTAGAGCTAGAAATAGCAAGTTAAAA |
| 12667 | 18566 | Pdcd1 | ATATATCTTGTGGAAAGGACGAAACACCG | GACACACGGCGCAATGACAG | GTTTTAGAGCTAGAAATAGCAAGTTAAAA | ATATATCTTGTGGAAAGGACGAAACACCGGACACACGGCGCAATGACAGGTTTTAGAGCTAGAAATAGCAAGTTAAAA |
| 12668 | 18566 | Pdcd1 | ATATATCTTGTGGAAAGGACGAAACACCG | CAGCTTGTCCAACTGGTCGG | GTTTTAGAGCTAGAAATAGCAAGTTAAAA | ATATATCTTGTGGAAAGGACGAAACACCGCAGCTTGTCCAACTGGTCGGGTTTTAGAGCTAGAAATAGCAAGTTAAAA |
| 12669 | 18566 | Pdcd1 | ATATATCTTGTGGAAAGGACGAAACACCG | GCTCAAACCATTACAGAAGG | GTTTTAGAGCTAGAAATAGCAAGTTAAAA | ATATATCTTGTGGAAAGGACGAAACACCGGCTCAAACCATTACAGAAGGGTTTTAGAGCTAGAAATAGCAAGTTAAAA |
| 3065 | 12778 | Ackr3 | ATATATCTTGTGGAAAGGACGAAACACCG | ACACCAAGATGCATACAACA | GTTTTAGAGCTAGAAATAGCAAGTTAAAA | ATATATCTTGTGGAAAGGACGAAACACCGACACCAAGATGCATACAACAGTTTTAGAGCTAGAAATAGCAAGTTAAAA |
| 3066 | 12778 | Ackr3 | ATATATCTTGTGGAAAGGACGAAACACCG | GCATAACCAGTGGCCCATGG | GTTTTAGAGCTAGAAATAGCAAGTTAAAA | ATATATCTTGTGGAAAGGACGAAACACCGGCATAACCAGTGGCCCATGGGTTTTAGAGCTAGAAATAGCAAGTTAAAA |
| 3067 | 12778 | Ackr3 | ATATATCTTGTGGAAAGGACGAAACACCG | GGTCCACGCTCATGCAGGCG | GTTTTAGAGCTAGAAATAGCAAGTTAAAA | ATATATCTTGTGGAAAGGACGAAACACCGGGTCCACGCTCATGCAGGCGGTTTTAGAGCTAGAAATAGCAAGTTAAAA |
| 3068 | 12778 | Ackr3 | ATATATCTTGTGGAAAGGACGAAACACCG | CCAACAATGAGACCTACTGC | GTTTTAGAGCTAGAAATAGCAAGTTAAAA | ATATATCTTGTGGAAAGGACGAAACACCGCCAACAATGAGACCTACTGCGTTTTAGAGCTAGAAATAGCAAGTTAAAA |
| | 110083 | DNAH12-1 | ATATATCTTGTGGAAAGGACGAAACACCG | AAAAGAATCACGTCGCATGG | GTTTTAGAGCTAGAAATAGCAAGTTAAAA | ATATATCTTGTGGAAAGGACGAAACACCGAAAAGAATCACGTCGCATGGGTTTTAGAGCTAGAAATAGCAAGTTAAAA |
| | 110083 | DNAH12-1 | ATATATCTTGTGGAAAGGACGAAACACCG | GAAATGATGATTGTTCGACA | GTTTTAGAGCTAGAAATAGCAAGTTAAAA | ATATATCTTGTGGAAAGGACGAAACACCGGAAATGATGATTGTTCGACAGTTTTAGAGCTAGAAATAGCAAGTTAAAA |
| | 110083 | DNAH12-2 | ATATATCTTGTGGAAAGGACGAAACACCG | CTGTAACATCTGTCAGTTAA | GTTTTAGAGCTAGAAATAGCAAGTTAAAA | ATATATCTTGTGGAAAGGACGAAACACCGCTGTAACATCTGTCAGTTAAGTTTTAGAGCTAGAAATAGCAAGTTAAAA |
| | 110083 | DNAH12-2 | ATATATCTTGTGGAAAGGACGAAACACCG | GGGCTAGACTATCTAGCAAT | GTTTTAGAGCTAGAAATAGCAAGTTAAAA | ATATATCTTGTGGAAAGGACGAAACACCGGGGCTAGACTATCTAGCAATGTTTTAGAGCTAGAAATAGCAAGTTAAAA |
| 5609 | 14198 | Fhit | ATATATCTTGTGGAAAGGACGAAACACCG | AACGGGTTTCCTATTCACCA | GTTTTAGAGCTAGAAATAGCAAGTTAAAA | ATATATCTTGTGGAAAGGACGAAACACCGAACGGGTTTCCTATTCACCAGTTTTAGAGCTAGAAATAGCAAGTTAAAA |
| 5610 | 14198 | Fhit | ATATATCTTGTGGAAAGGACGAAACACCG | CTTCATCAGGATGTAGGTCA | GTTTTAGAGCTAGAAATAGCAAGTTAAAA | ATATATCTTGTGGAAAGGACGAAACACCGCTTCATCAGGATGTAGGTCAGTTTTAGAGCTAGAAATAGCAAGTTAAAA |
| 5611 | 14198 | Fhit | ATATATCTTGTGGAAAGGACGAAACACCG | AAGCGCTCTACTGGCCTCAG | GTTTTAGAGCTAGAAATAGCAAGTTAAAA | ATATATCTTGTGGAAAGGACGAAACACCGAAGCGCTCTACTGGCCTCAGGTTTTAGAGCTAGAAATAGCAAGTTAAAA |
| 5612 | 14198 | Fhit | ATATATCTTGTGGAAAGGACGAAACACCG | CCAGAGAGTTGGGACAGTGG | GTTTTAGAGCTAGAAATAGCAAGTTAAAA | ATATATCTTGTGGAAAGGACGAAACACCGCCAGAGAGTTGGGACAGTGGGTTTTAGAGCTAGAAATAGCAAGTTAAAA |
| 8685 | 16331 | Inpp5d | ATATATCTTGTGGAAAGGACGAAACACCG | TGGGGATCTCAACTACCGCG | GTTTTAGAGCTAGAAATAGCAAGTTAAAA | ATATATCTTGTGGAAAGGACGAAACACCGTGGGGATCTCAACTACCGCGGTTTTAGAGCTAGAAATAGCAAGTTAAAA |
| 8686 | 16331 | Inpp5d | ATATATCTTGTGGAAAGGACGAAACACCG | CCACTATGCGAATGTTCCAG | GTTTTAGAGCTAGAAATAGCAAGTTAAAA | ATATATCTTGTGGAAAGGACGAAACACCGCCACTATGCGAATGTTCCAGGTTTTAGAGCTAGAAATAGCAAGTTAAAA |
| 8687 | 16331 | Inpp5d | ATATATCTTGTGGAAAGGACGAAACACCG | AGAGGCCAGTCCCATCACCA | GTTTTAGAGCTAGAAATAGCAAGTTAAAA | ATATATCTTGTGGAAAGGACGAAACACCGAGAGGCCAGTCCCATCACCAGTTTTAGAGCTAGAAATAGCAAGTTAAAA |
| 8688 | 16331 | Inpp5d | ATATATCTTGTGGAAAGGACGAAACACCG | CATCCGAAGGTGTCCCCATG | GTTTTAGAGCTAGAAATAGCAAGTTAAAA | ATATATCTTGTGGAAAGGACGAAACACCGCATCCGAAGGTGTCCCCATGGTTTTAGAGCTAGAAATAGCAAGTTAAAA |
| 35406 | 70122 | Mllt3 | ATATATCTTGTGGAAAGGACGAAACACCG | AACTTTCAATAACCCCACGG | GTTTTAGAGCTAGAAATAGCAAGTTAAAA | ATATATCTTGTGGAAAGGACGAAACACCGAACTTTCAATAACCCCACGGGTTTTAGAGCTAGAAATAGCAAGTTAAAA |
| 35407 | 70122 | Mllt3 | ATATATCTTGTGGAAAGGACGAAACACCG | AAGGAAAAAGAGTAGCTCGG | GTTTTAGAGCTAGAAATAGCAAGTTAAAA | ATATATCTTGTGGAAAGGACGAAACACCGAAGGAAAAAGAGTAGCTCGGGTTTTAGAGCTAGAAATAGCAAGTTAAAA |
| 35408 | 70122 | Mllt3 | ATATATCTTGTGGAAAGGACGAAACACCG | TCCACGATGTCATCAAACGG | GTTTTAGAGCTAGAAATAGCAAGTTAAAA | ATATATCTTGTGGAAAGGACGAAACACCGTCCACGATGTCATCAAACGGGTTTTAGAGCTAGAAATAGCAAGTTAAAA |
| 35409 | 70122 | Mllt3 | ATATATCTTGTGGAAAGGACGAAACACCG | ACTTACTCACCGTCACCAGT | GTTTTAGAGCTAGAAATAGCAAGTTAAAA | ATATATCTTGTGGAAAGGACGAAACACCGACTTACTCACCGTCACCAGTGTTTTAGAGCTAGAAATAGCAAGTTAAAA |
| 15134 | 20190 | Ryr1 | ATATATCTTGTGGAAAGGACGAAACACCG | CTTCAAACTCGAAGTACCAG | GTTTTAGAGCTAGAAATAGCAAGTTAAAA | ATATATCTTGTGGAAAGGACGAAACACCGCTTCAAACTCGAAGTACCAGGTTTTAGAGCTAGAAATAGCAAGTTAAAA |
| 15135 | 20190 | Ryr1 | ATATATCTTGTGGAAAGGACGAAACACCG | GGGACAACGTAACCACGCTG | GTTTTAGAGCTAGAAATAGCAAGTTAAAA | ATATATCTTGTGGAAAGGACGAAACACCGGGGACAACGTAACCACGCTGGTTTTAGAGCTAGAAATAGCAAGTTAAAA |
| 15136 | 20190 | Ryr1 | ATATATCTTGTGGAAAGGACGAAACACCG | GCGATTGCCACGGATCAGAG | GTTTTAGAGCTAGAAATAGCAAGTTAAAA | ATATATCTTGTGGAAAGGACGAAACACCGGCGATTGCCACGGATCAGAGGTTTTAGAGCTAGAAATAGCAAGTTAAAA |
| 15137 | 20190 | Ryr1 | ATATATCTTGTGGAAAGGACGAAACACCG | CATCAAAGAGTATCGCAGGG | GTTTTAGAGCTAGAAATAGCAAGTTAAAA | ATATATCTTGTGGAAAGGACGAAACACCGCATCAAAGAGTATCGCAGGGGTTTTAGAGCTAGAAATAGCAAGTTAAAA |
| 60580 | 235587 | Parp3 | ATATATCTTGTGGAAAGGACGAAACACCG | TGATAATCGGGTCATCCGTG | GTTTTAGAGCTAGAAATAGCAAGTTAAAA | ATATATCTTGTGGAAAGGACGAAACACCGTGATAATCGGGTCATCCGTGGTTTTAGAGCTAGAAATAGCAAGTTAAAA |
| 60581 | 235587 | Parp3 | ATATATCTTGTGGAAAGGACGAAACACCG | GCTGGTAGTCTCGATCCAGT | GTTTTAGAGCTAGAAATAGCAAGTTAAAA | ATATATCTTGTGGAAAGGACGAAACACCGGCTGGTAGTCTCGATCCAGTGTTTTAGAGCTAGAAATAGCAAGTTAAAA |
| 60582 | 235587 | Parp3 | ATATATCTTGTGGAAAGGACGAAACACCG | GAAGACTAAAAACAAATGGG | GTTTTAGAGCTAGAAATAGCAAGTTAAAA | ATATATCTTGTGGAAAGGACGAAACACCGGAAGACTAAAAACAAATGGGGTTTTAGAGCTAGAAATAGCAAGTTAAAA |
| 60583 | 235587 | Parp3 | ATATATCTTGTGGAAAGGACGAAACACCG | ACCCTGAACCAGACCAACAT | GTTTTAGAGCTAGAAATAGCAAGTTAAAA | ATATATCTTGTGGAAAGGACGAAACACCGACCCTGAACCAGACCAACATGTTTTAGAGCTAGAAATAGCAAGTTAAAA |
| 14830 | 19882 | Mst1r | ATATATCTTGTGGAAAGGACGAAACACCG | AAGTATCAGACTTTAGACGA | GTTTTAGAGCTAGAAATAGCAAGTTAAAA | ATATATCTTGTGGAAAGGACGAAACACCGAAGTATCAGACTTTAGACGAGTTTTAGAGCTAGAAATAGCAAGTTAAAA |
| 14831 | 19882 | Mst1r | ATATATCTTGTGGAAAGGACGAAACACCG | GGGAACACACCAGATCACCG | GTTTTAGAGCTAGAAATAGCAAGTTAAAA | ATATATCTTGTGGAAAGGACGAAACACCGGGGAACACACCAGATCACCGGTTTTAGAGCTAGAAATAGCAAGTTAAAA |
| 14832 | 19882 | Mst1r | ATATATCTTGTGGAAAGGACGAAACACCG | ATGTGGTCCGAAACCCTCAG | GTTTTAGAGCTAGAAATAGCAAGTTAAAA | ATATATCTTGTGGAAAGGACGAAACACCGATGTGGTCCGAAACCCTCAGGTTTTAGAGCTAGAAATAGCAAGTTAAAA |
| 14833 | 19882 | Mst1r | ATATATCTTGTGGAAAGGACGAAACACCG | AAGACGTGTCACATGCAGTG | GTTTTAGAGCTAGAAATAGCAAGTTAAAA | ATATATCTTGTGGAAAGGACGAAACACCGAAGACGTGTCACATGCAGTGGTTTTAGAGCTAGAAATAGCAAGTTAAAA |
| 7561 | 15235 | Mst1 | ATATATCTTGTGGAAAGGACGAAACACCG | CGGTGAGGATTACCGTGGCG | GTTTTAGAGCTAGAAATAGCAAGTTAAAA | ATATATCTTGTGGAAAGGACGAAACACCGCGGTGAGGATTACCGTGGCGGTTTTAGAGCTAGAAATAGCAAGTTAAAA |
| 7562 | 15235 | Mst1 | ATATATCTTGTGGAAAGGACGAAACACCG | GGCTCAGGTGAACAGTATCG | GTTTTAGAGCTAGAAATAGCAAGTTAAAA | ATATATCTTGTGGAAAGGACGAAACACCGGGCTCAGGTGAACAGTATCGGTTTTAGAGCTAGAAATAGCAAGTTAAAA |
| 7563 | 15235 | Mst1 | ATATATCTTGTGGAAAGGACGAAACACCG | AAGTCACAGCGGCGCAACAA | GTTTTAGAGCTAGAAATAGCAAGTTAAAA | ATATATCTTGTGGAAAGGACGAAACACCGAAGTCACAGCGGCGCAACAAGTTTTAGAGCTAGAAATAGCAAGTTAAAA |
| 7564 | 15235 | Mst1 | ATATATCTTGTGGAAAGGACGAAACACCG | GTAATCCGGACGGATCTGAG | GTTTTAGAGCTAGAAATAGCAAGTTAAAA | ATATATCTTGTGGAAAGGACGAAACACCGGTAATCCGGACGGATCTGAGGTTTTAGAGCTAGAAATAGCAAGTTAAAA |
| 43128 | 76804 | Kdm4c | ATATATCTTGTGGAAAGGACGAAACACCG | GTGCACTTGTCGGAATGACA | GTTTTAGAGCTAGAAATAGCAAGTTAAAA | ATATATCTTGTGGAAAGGACGAAACACCGGTGCACTTGTCGGAATGACAGTTTTAGAGCTAGAAATAGCAAGTTAAAA |
| 43129 | 76804 | Kdm4c | ATATATCTTGTGGAAAGGACGAAACACCG | CTGGCCGGAGGCTTACCAAG | GTTTTAGAGCTAGAAATAGCAAGTTAAAA | ATATATCTTGTGGAAAGGACGAAACACCGCTGGCCGGAGGCTTACCAAGGTTTTAGAGCTAGAAATAGCAAGTTAAAA |
| 43130 | 76804 | Kdm4c | ATATATCTTGTGGAAAGGACGAAACACCG | TTTGGATACCAGGATACAAG | GTTTTAGAGCTAGAAATAGCAAGTTAAAA | ATATATCTTGTGGAAAGGACGAAACACCGTTTGGATACCAGGATACAAGGTTTTAGAGCTAGAAATAGCAAGTTAAAA |
| 43131 | 76804 | Kdm4c | ATATATCTTGTGGAAAGGACGAAACACCG | TGTGGAAAACAACATTTGCG | GTTTTAGAGCTAGAAATAGCAAGTTAAAA | ATATATCTTGTGGAAAGGACGAAACACCGTGTGGAAAACAACATTTGCGGTTTTAGAGCTAGAAATAGCAAGTTAAAA |
| 47196 | 101739 | Psip1 | ATATATCTTGTGGAAAGGACGAAACACCG | AGAGAGGACGACCTGCAGGT | GTTTTAGAGCTAGAAATAGCAAGTTAAAA | ATATATCTTGTGGAAAGGACGAAACACCGAGAGAGGACGACCTGCAGGTGTTTTAGAGCTAGAAATAGCAAGTTAAAA |
| 47197 | 101739 | Psip1 | ATATATCTTGTGGAAAGGACGAAACACCG | GAAAAATTTAGCTAAACCAG | GTTTTAGAGCTAGAAATAGCAAGTTAAAA | ATATATCTTGTGGAAAGGACGAAACACCGGAAAAATTTAGCTAAACCAGGTTTTAGAGCTAGAAATAGCAAGTTAAAA |
| 47198 | 101739 | Psip1 | ATATATCTTGTGGAAAGGACGAAACACCG | ACTCCAAAAGCTGCCAGGCG | GTTTTAGAGCTAGAAATAGCAAGTTAAAA | ATATATCTTGTGGAAAGGACGAAACACCGACTCCAAAAGCTGCCAGGCGGTTTTAGAGCTAGAAATAGCAAGTTAAAA |
| 47199 | 101739 | Psip1 | ATATATCTTGTGGAAAGGACGAAACACCG | AAAAGAGACTAACGTTTCAA | GTTTTAGAGCTAGAAATAGCAAGTTAAAA | ATATATCTTGTGGAAAGGACGAAACACCGAAAAGAGACTAACGTTTCAAGTTTTAGAGCTAGAAATAGCAAGTTAAAA |
| 29486 | 66923 | Pbrm1 | ATATATCTTGTGGAAAGGACGAAACACCG | AAAACACTTGCATAACGATG | GTTTTAGAGCTAGAAATAGCAAGTTAAAA | ATATATCTTGTGGAAAGGACGAAACACCGAAAACACTTGCATAACGATGGTTTTAGAGCTAGAAATAGCAAGTTAAAA |
| 29487 | 66923 | Pbrm1 | ATATATCTTGTGGAAAGGACGAAACACCG | CAATGCCAGGCACTACAATG | GTTTTAGAGCTAGAAATAGCAAGTTAAAA | ATATATCTTGTGGAAAGGACGAAACACCGCAATGCCAGGCACTACAATGGTTTTAGAGCTAGAAATAGCAAGTTAAAA |
| 29488 | 66923 | Pbrm1 | ATATATCTTGTGGAAAGGACGAAACACCG | AATAAAAGAGCAGTCCAAGG | GTTTTAGAGCTAGAAATAGCAAGTTAAAA | ATATATCTTGTGGAAAGGACGAAACACCGAATAAAAGAGCAGTCCAAGGGTTTTAGAGCTAGAAATAGCAAGTTAAAA |
| 29489 | 66923 | Pbrm1 | ATATATCTTGTGGAAAGGACGAAACACCG | ACAAGCAGCTTTATACTCAG | GTTTTAGAGCTAGAAATAGCAAGTTAAAA | ATATATCTTGTGGAAAGGACGAAACACCGACAAGCAGCTTTATACTCAGGTTTTAGAGCTAGAAATAGCAAGTTAAAA |
| 47816 | 104416 | Bap1 | ATATATCTTGTGGAAAGGACGAAACACCG | CCACCAACGTAGAAACCTTG | GTTTTAGAGCTAGAAATAGCAAGTTAAAA | ATATATCTTGTGGAAAGGACGAAACACCGCCACCAACGTAGAAACCTTGGTTTTAGAGCTAGAAATAGCAAGTTAAAA |
| 47817 | 104416 | Bap1 | ATATATCTTGTGGAAAGGACGAAACACCG | GCAGGATTAAGTATGAGACC | GTTTTAGAGCTAGAAATAGCAAGTTAAAA | ATATATCTTGTGGAAAGGACGAAACACCGGCAGGATTAAGTATGAGACCGTTTTAGAGCTAGAAATAGCAAGTTAAAA |
| 47818 | 104416 | Bap1 | ATATATCTTGTGGAAAGGACGAAACACCG | ACCTGTCTGAGTGCACTCAG | GTTTTAGAGCTAGAAATAGCAAGTTAAAA | ATATATCTTGTGGAAAGGACGAAACACCGACCTGTCTGAGTGCACTCAGGTTTTAGAGCTAGAAATAGCAAGTTAAAA |
| 47819 | 104416 | Bap1 | ATATATCTTGTGGAAAGGACGAAACACCG | TCAGCTATGTGCCTATCACA | GTTTTAGAGCTAGAAATAGCAAGTTAAAA | ATATATCTTGTGGAAAGGACGAAACACCGTCAGCTATGTGCCTATCACAGTTTTAGAGCTAGAAATAGCAAGTTAAAA |
| 26086 | 58205 | Pdcd1lg2 | ATATATCTTGTGGAAAGGACGAAACACCG | CCCCGCAGATGACCAGGCAA | GTTTTAGAGCTAGAAATAGCAAGTTAAAA | ATATATCTTGTGGAAAGGACGAAACACCGCCCCGCAGATGACCAGGCAAGTTTTAGAGCTAGAAATAGCAAGTTAAAA |
| 26087 | 58205 | Pdcd1lg2 | ATATATCTTGTGGAAAGGACGAAACACCG | GATAGACACTAGGATCCTGG | GTTTTAGAGCTAGAAATAGCAAGTTAAAA | ATATATCTTGTGGAAAGGACGAAACACCGGATAGACACTAGGATCCTGGGTTTTAGAGCTAGAAATAGCAAGTTAAAA |
| 26088 | 58205 | Pdcd1lg2 | ATATATCTTGTGGAAAGGACGAAACACCG | CCAGTTCAGTGCATTCTCTG | GTTTTAGAGCTAGAAATAGCAAGTTAAAA | ATATATCTTGTGGAAAGGACGAAACACCGCCAGTTCAGTGCATTCTCTGGTTTTAGAGCTAGAAATAGCAAGTTAAAA |
| 26089 | 58205 | Pdcd1lg2 | ATATATCTTGTGGAAAGGACGAAACACCG | CTAGGGGATAACCTCTAGCC | GTTTTAGAGCTAGAAATAGCAAGTTAAAA | ATATATCTTGTGGAAAGGACGAAACACCGCTAGGGGATAACCTCTAGCCGTTTTAGAGCTAGAAATAGCAAGTTAAAA |
| 8933 | 16452 | Jak2 | ATATATCTTGTGGAAAGGACGAAACACCG | CGGGTATTACAGACTAACTG | GTTTTAGAGCTAGAAATAGCAAGTTAAAA | ATATATCTTGTGGAAAGGACGAAACACCGCGGGTATTACAGACTAACTGGTTTTAGAGCTAGAAATAGCAAGTTAAAA |
| 8934 | 16452 | Jak2 | ATATATCTTGTGGAAAGGACGAAACACCG | AAGTCCTAGATAAAGCACAT | GTTTTAGAGCTAGAAATAGCAAGTTAAAA | ATATATCTTGTGGAAAGGACGAAACACCGAAGTCCTAGATAAAGCACATGTTTTAGAGCTAGAAATAGCAAGTTAAAA |
| 8935 | 16452 | Jak2 | ATATATCTTGTGGAAAGGACGAAACACCG | ACAGATACGGAGTGTCCCGT | GTTTTAGAGCTAGAAATAGCAAGTTAAAA | ATATATCTTGTGGAAAGGACGAAACACCGACAGATACGGAGTGTCCCGTGTTTTAGAGCTAGAAATAGCAAGTTAAAA |
| 8936 | 16452 | Jak2 | ATATATCTTGTGGAAAGGACGAAACACCG | TCTTCAGGAGAGAATACCAT | GTTTTAGAGCTAGAAATAGCAAGTTAAAA | ATATATCTTGTGGAAAGGACGAAACACCGTCTTCAGGAGAGAATACCATGTTTTAGAGCTAGAAATAGCAAGTTAAAA |
| 8737 | 16367 | Irs1 | ATATATCTTGTGGAAAGGACGAAACACCG | ACCCACGCCCATCGGCATCG | GTTTTAGAGCTAGAAATAGCAAGTTAAAA | ATATATCTTGTGGAAAGGACGAAACACCGACCCACGCCCATCGGCATCGGTTTTAGAGCTAGAAATAGCAAGTTAAAA |
| 8738 | 16367 | Irs1 | ATATATCTTGTGGAAAGGACGAAACACCG | ATGCACCACTTGGAACGTCG | GTTTTAGAGCTAGAAATAGCAAGTTAAAA | ATATATCTTGTGGAAAGGACGAAACACCGATGCACCACTTGGAACGTCGGTTTTAGAGCTAGAAATAGCAAGTTAAAA |
| 8739 | 16367 | Irs1 | ATATATCTTGTGGAAAGGACGAAACACCG | GGGAATCTGGGGTGACACTG | GTTTTAGAGCTAGAAATAGCAAGTTAAAA | ATATATCTTGTGGAAAGGACGAAACACCGGGGAATCTGGGGTGACACTGGTTTTAGAGCTAGAAATAGCAAGTTAAAA |
| 8740 | 16367 | Irs1 | ATATATCTTGTGGAAAGGACGAAACACCG | GGAGGACTTGAGCTATGACA | GTTTTAGAGCTAGAAATAGCAAGTTAAAA | ATATATCTTGTGGAAAGGACGAAACACCGGGAGGACTTGAGCTATGACAGTTTTAGAGCTAGAAATAGCAAGTTAAAA |
| 26694 | 60533 | Cd274 | ATATATCTTGTGGAAAGGACGAAACACCG | TGCTGCATAATCAGCTACGG | GTTTTAGAGCTAGAAATAGCAAGTTAAAA | ATATATCTTGTGGAAAGGACGAAACACCGTGCTGCATAATCAGCTACGGGTTTTAGAGCTAGAAATAGCAAGTTAAAA |
| 26695 | 60533 | Cd274 | ATATATCTTGTGGAAAGGACGAAACACCG | TCCAAAGGACTTGTACGTGG | GTTTTAGAGCTAGAAATAGCAAGTTAAAA | ATATATCTTGTGGAAAGGACGAAACACCGTCCAAAGGACTTGTACGTGGGTTTTAGAGCTAGAAATAGCAAGTTAAAA |
| 26696 | 60533 | Cd274 | ATATATCTTGTGGAAAGGACGAAACACCG | CTCAGCACAGCAACTTCAGG | GTTTTAGAGCTAGAAATAGCAAGTTAAAA | ATATATCTTGTGGAAAGGACGAAACACCGCTCAGCACAGCAACTTCAGGGTTTTAGAGCTAGAAATAGCAAGTTAAAA |
| 26697 | 60533 | Cd274 | ATATATCTTGTGGAAAGGACGAAACACCG | GCTTGCGTTAGTGGTGTACT | GTTTTAGAGCTAGAAATAGCAAGTTAAAA | ATATATCTTGTGGAAAGGACGAAACACCGGCTTGCGTTAGTGGTGTACTGTTTTAGAGCTAGAAATAGCAAGTTAAAA |
| 12026 | 18223 | Numbl | ATATATCTTGTGGAAAGGACGAAACACCG | CTATCCTATGTCACTAGACG | GTTTTAGAGCTAGAAATAGCAAGTTAAAA | ATATATCTTGTGGAAAGGACGAAACACCGCTATCCTATGTCACTAGACGGTTTTAGAGCTAGAAATAGCAAGTTAAAA |
| 12027 | 18223 | Numbl | ATATATCTTGTGGAAAGGACGAAACACCG | ATGGCAGCTCATTCAACCGT | GTTTTAGAGCTAGAAATAGCAAGTTAAAA | ATATATCTTGTGGAAAGGACGAAACACCGATGGCAGCTCATTCAACCGTGTTTTAGAGCTAGAAATAGCAAGTTAAAA |
| 12028 | 18223 | Numbl | ATATATCTTGTGGAAAGGACGAAACACCG | TTCTCCTACATATGCCGCGA | GTTTTAGAGCTAGAAATAGCAAGTTAAAA | ATATATCTTGTGGAAAGGACGAAACACCGTTCTCCTACATATGCCGCGAGTTTTAGAGCTAGAAATAGCAAGTTAAAA |
| 12029 | 18223 | Numbl | ATATATCTTGTGGAAAGGACGAAACACCG | CCCGTGCACTCACCCTAACA | GTTTTAGAGCTAGAAATAGCAAGTTAAAA | ATATATCTTGTGGAAAGGACGAAACACCGCCCGTGCACTCACCCTAACAGTTTTAGAGCTAGAAATAGCAAGTTAAAA |
| 3409 | 12914 | Crebbp | ATATATCTTGTGGAAAGGACGAAACACCG | CATACTCGAGATATCCGATG | GTTTTAGAGCTAGAAATAGCAAGTTAAAA | ATATATCTTGTGGAAAGGACGAAACACCGCATACTCGAGATATCCGATGGTTTTAGAGCTAGAAATAGCAAGTTAAAA |
| 3410 | 12914 | Crebbp | ATATATCTTGTGGAAAGGACGAAACACCG | AGCTCTGAAAGATCGCCGCA | GTTTTAGAGCTAGAAATAGCAAGTTAAAA | ATATATCTTGTGGAAAGGACGAAACACCGAGCTCTGAAAGATCGCCGCAGTTTTAGAGCTAGAAATAGCAAGTTAAAA |
| 3411 | 12914 | Crebbp | ATATATCTTGTGGAAAGGACGAAACACCG | TAATGAATCAGGCTCAACAA | GTTTTAGAGCTAGAAATAGCAAGTTAAAA | ATATATCTTGTGGAAAGGACGAAACACCGTAATGAATCAGGCTCAACAAGTTTTAGAGCTAGAAATAGCAAGTTAAAA |
| 3412 | 12914 | Crebbp | ATATATCTTGTGGAAAGGACGAAACACCG | TGAACCTACTGAATCCAAGG | GTTTTAGAGCTAGAAATAGCAAGTTAAAA | ATATATCTTGTGGAAAGGACGAAACACCGTGAACCTACTGAATCCAAGGGTTTTAGAGCTAGAAATAGCAAGTTAAAA |
| 14934 | 19989 | Rpl7 | ATATATCTTGTGGAAAGGACGAAACACCG | GAAGGCAAAGCACTATCACA | GTTTTAGAGCTAGAAATAGCAAGTTAAAA | ATATATCTTGTGGAAAGGACGAAACACCGGAAGGCAAAGCACTATCACAGTTTTAGAGCTAGAAATAGCAAGTTAAAA |
| 14935 | 19989 | Rpl7 | ATATATCTTGTGGAAAGGACGAAACACCG | CCTTTGTCATCAGAATTCGA | GTTTTAGAGCTAGAAATAGCAAGTTAAAA | ATATATCTTGTGGAAAGGACGAAACACCGCCTTTGTCATCAGAATTCGAGTTTTAGAGCTAGAAATAGCAAGTTAAAA |
| 14936 | 19989 | Rpl7 | ATATATCTTGTGGAAAGGACGAAACACCG | TGGAGCCATACATTGCATGG | GTTTTAGAGCTAGAAATAGCAAGTTAAAA | ATATATCTTGTGGAAAGGACGAAACACCGTGGAGCCATACATTGCATGGGTTTTAGAGCTAGAAATAGCAAGTTAAAA |
| 14937 | 19989 | Rpl7 | ATATATCTTGTGGAAAGGACGAAACACCG | TTAAGAAAAAGGTTCCTGCT | GTTTTAGAGCTAGAAATAGCAAGTTAAAA | ATATATCTTGTGGAAAGGACGAAACACCGTTAAGAAAAAGGTTCCTGCTGTTTTAGAGCTAGAAATAGCAAGTTAAAA |
| 15006 | 20085 | Rps19 | ATATATCTTGTGGAAAGGACGAAACACCG | GCACGGCACCTGTACCTCCG | GTTTTAGAGCTAGAAATAGCAAGTTAAAA | ATATATCTTGTGGAAAGGACGAAACACCGGCACGGCACCTGTACCTCCGGTTTTAGAGCTAGAAATAGCAAGTTAAAA |
| 15007 | 20085 | Rps19 | ATATATCTTGTGGAAAGGACGAAACACCG | AACTGGTTCTACACACGAGC | GTTTTAGAGCTAGAAATAGCAAGTTAAAA | ATATATCTTGTGGAAAGGACGAAACACCGAACTGGTTCTACACACGAGCGTTTTAGAGCTAGAAATAGCAAGTTAAAA |
| 15008 | 20085 | Rps19 | ATATATCTTGTGGAAAGGACGAAACACCG | TCCATGACCAAGATCTATGG | GTTTTAGAGCTAGAAATAGCAAGTTAAAA | ATATATCTTGTGGAAAGGACGAAACACCGTCCATGACCAAGATCTATGGGTTTTAGAGCTAGAAATAGCAAGTTAAAA |
| 15009 | 20085 | Rps19 | ATATATCTTGTGGAAAGGACGAAACACCG | AGACCCAGCCATTTCAGCAG | GTTTTAGAGCTAGAAATAGCAAGTTAAAA | ATATATCTTGTGGAAAGGACGAAACACCGAGACCCAGCCATTTCAGCAGGTTTTAGAGCTAGAAATAGCAAGTTAAAA |
| 14890 | 19934 | Rpl22 | ATATATCTTGTGGAAAGGACGAAACACCG | AAAAAGCTTGTGGCGAAGGG | GTTTTAGAGCTAGAAATAGCAAGTTAAAA | ATATATCTTGTGGAAAGGACGAAACACCGAAAAAGCTTGTGGCGAAGGGGTTTTAGAGCTAGAAATAGCAAGTTAAAA |
| 14891 | 19934 | Rpl22 | ATATATCTTGTGGAAAGGACGAAACACCG | CAAGATCACTGTCACTTCAG | GTTTTAGAGCTAGAAATAGCAAGTTAAAA | ATATATCTTGTGGAAAGGACGAAACACCGCAAGATCACTGTCACTTCAGGTTTTAGAGCTAGAAATAGCAAGTTAAAA |
| 14892 | 19934 | Rpl22 | ATATATCTTGTGGAAAGGACGAAACACCG | TGCTACGAACCTTTCTGCTC | GTTTTAGAGCTAGAAATAGCAAGTTAAAA | ATATATCTTGTGGAAAGGACGAAACACCGTGCTACGAACCTTTCTGCTCGTTTTAGAGCTAGAAATAGCAAGTTAAAA |
| 14893 | 19934 | Rpl22 | ATATATCTTGTGGAAAGGACGAAACACCG | CAGTTAAGTTCTCTAAGAAG | GTTTTAGAGCTAGAAATAGCAAGTTAAAA | ATATATCTTGTGGAAAGGACGAAACACCGCAGTTAAGTTCTCTAAGAAGGTTTTAGAGCTAGAAATAGCAAGTTAAAA |
| 15002 | 20084 | Rps18 | ATATATCTTGTGGAAAGGACGAAACACCG | TTTGACCTCAGGGCGTGGGG | GTTTTAGAGCTAGAAATAGCAAGTTAAAA | ATATATCTTGTGGAAAGGACGAAACACCGTTTGACCTCAGGGCGTGGGGGTTTTAGAGCTAGAAATAGCAAGTTAAAA |
| 15003 | 20084 | Rps18 | ATATATCTTGTGGAAAGGACGAAACACCG | AGACATCGACCTCACCAAGA | GTTTTAGAGCTAGAAATAGCAAGTTAAAA | ATATATCTTGTGGAAAGGACGAAACACCGAGACATCGACCTCACCAAGAGTTTTAGAGCTAGAAATAGCAAGTTAAAA |
| 15004 | 20084 | Rps18 | ATATATCTTGTGGAAAGGACGAAACACCG | GACAGTACAAGATCCCAGAC | GTTTTAGAGCTAGAAATAGCAAGTTAAAA | ATATATCTTGTGGAAAGGACGAAACACCGGACAGTACAAGATCCCAGACGTTTTAGAGCTAGAAATAGCAAGTTAAAA |
| 15005 | 20084 | Rps18 | ATATATCTTGTGGAAAGGACGAAACACCG | AAAATCTAGGTTCTGGCCAA | GTTTTAGAGCTAGAAATAGCAAGTTAAAA | ATATATCTTGTGGAAAGGACGAAACACCGAAAATCTAGGTTCTGGCCAAGTTTTAGAGCTAGAAATAGCAAGTTAAAA |
| 20826 | 27207 | Rps11 | ATATATCTTGTGGAAAGGACGAAACACCG | CGATATTCTTGTAGTACCGA | GTTTTAGAGCTAGAAATAGCAAGTTAAAA | ATATATCTTGTGGAAAGGACGAAACACCGCGATATTCTTGTAGTACCGAGTTTTAGAGCTAGAAATAGCAAGTTAAAA |
| 20827 | 27207 | Rps11 | ATATATCTTGTGGAAAGGACGAAACACCG | AGGCTTCAAGACGCCTAAAG | GTTTTAGAGCTAGAAATAGCAAGTTAAAA | ATATATCTTGTGGAAAGGACGAAACACCGAGGCTTCAAGACGCCTAAAGGTTTTAGAGCTAGAAATAGCAAGTTAAAA |
| 20828 | 27207 | Rps11 | ATATATCTTGTGGAAAGGACGAAACACCG | ATAGTCCCGGCGGATGACAA | GTTTTAGAGCTAGAAATAGCAAGTTAAAA | ATATATCTTGTGGAAAGGACGAAACACCGATAGTCCCGGCGGATGACAAGTTTTAGAGCTAGAAATAGCAAGTTAAAA |
| 20829 | 27207 | Rps11 | ATATATCTTGTGGAAAGGACGAAACACCG | CAACATCCCACTCACCAGAC | GTTTTAGAGCTAGAAATAGCAAGTTAAAA | ATATATCTTGTGGAAAGGACGAAACACCGCAACATCCCACTCACCAGACGTTTTAGAGCTAGAAATAGCAAGTTAAAA |
| 83078 | NonTarge | MGLibA_66406 | ATATATCTTGTGGAAAGGACGAAACACCG | GCGAGGTATTCGGCTCCGCG | GTTTTAGAGCTAGAAATAGCAAGTTAAAA | ATATATCTTGTGGAAAGGACGAAACACCGGCGAGGTATTCGGCTCCGCGGTTTTAGAGCTAGAAATAGCAAGTTAAAA |
| 83079 | NonTarge | MGLibA_66407 | ATATATCTTGTGGAAAGGACGAAACACCG | GCTTTCACGGAGGTTCGACG | GTTTTAGAGCTAGAAATAGCAAGTTAAAA | ATATATCTTGTGGAAAGGACGAAACACCGGCTTTCACGGAGGTTCGACGGTTTTAGAGCTAGAAATAGCAAGTTAAAA |
| 83080 | NonTarge | MGLibA_66408 | ATATATCTTGTGGAAAGGACGAAACACCG | ATGTTGCAGTTCGGCTCGAT | GTTTTAGAGCTAGAAATAGCAAGTTAAAA | ATATATCTTGTGGAAAGGACGAAACACCGATGTTGCAGTTCGGCTCGATGTTTTAGAGCTAGAAATAGCAAGTTAAAA |
| 83081 | NonTarge | MGLibA_66409 | ATATATCTTGTGGAAAGGACGAAACACCG | ACGTGTAAGGCGAACGCCTT | GTTTTAGAGCTAGAAATAGCAAGTTAAAA | ATATATCTTGTGGAAAGGACGAAACACCGACGTGTAAGGCGAACGCCTTGTTTTAGAGCTAGAAATAGCAAGTTAAAA |
| 83082 | NonTarge | MGLibA_66410 | ATATATCTTGTGGAAAGGACGAAACACCG | GACTCCGGGTACTAAATGTC | GTTTTAGAGCTAGAAATAGCAAGTTAAAA | ATATATCTTGTGGAAAGGACGAAACACCGGACTCCGGGTACTAAATGTCGTTTTAGAGCTAGAAATAGCAAGTTAAAA |
| 83083 | NonTarge | MGLibA_66411 | ATATATCTTGTGGAAAGGACGAAACACCG | CCGCGCCGTTAGGGAACGAG | GTTTTAGAGCTAGAAATAGCAAGTTAAAA | ATATATCTTGTGGAAAGGACGAAACACCGCCGCGCCGTTAGGGAACGAGGTTTTAGAGCTAGAAATAGCAAGTTAAAA |
| 83084 | NonTarge | MGLibA_66412 | ATATATCTTGTGGAAAGGACGAAACACCG | ATTGTTCGACCGTCTACGGG | GTTTTAGAGCTAGAAATAGCAAGTTAAAA | ATATATCTTGTGGAAAGGACGAAACACCGATTGTTCGACCGTCTACGGGGTTTTAGAGCTAGAAATAGCAAGTTAAAA |
| 83085 | NonTarge | MGLibA_66413 | ATATATCTTGTGGAAAGGACGAAACACCG | ACCCATCGGGTGCGATATGG | GTTTTAGAGCTAGAAATAGCAAGTTAAAA | ATATATCTTGTGGAAAGGACGAAACACCGACCCATCGGGTGCGATATGGGTTTTAGAGCTAGAAATAGCAAGTTAAAA |

Supplementary Table 6. Combined T-cell/MDSC panel.

| Location | Antibody | Clone | Source |
|---|---|---|---|
| | Live/dead | | ebio (65-0865-18) |
| | CD45 | 30-F11 | BioLegend |
| | CD4 | RM4-5 | BioLegend |
| | CD8 | 53-6.7 | BioLegend |
| | NKp46 | 29A1.4 | BioLegend |
| | PD-1 | J43 | ThermoFisher |
| | Tim-3 | 215008 | R&D Systems |
| | Lag-3 | C9B7W | BioLegend |
| | CD122 | TM-B1 | BD Biosciences |
| Surface | CD62L | MEL-14 | BioLegend |
| | MHCII | 2G9 | BD Biosciences |
| | KLRG1 | 2F1/KLRG1 | BioLegend |
| | F4/80 | T45-2342 | BD Biosciences |
| | CD11c | N418 | ThermoFisher |
| | CD11b | M1/70 | ThermoFisher |
| | CD274 (PD-L1) | 10F.9G2 | BD Biosciences |
| | Ly6C | HK1.4 | BioLegend |
| | Ly6G | 1A8 | BD Biosciences |
| | Granzyme A | GzA-3G8.5 | ThermoFisher |
| | FoxP3 | FKJ-16a | ThermoFisher |
| Intracellular | Ki-67 | B56 | BD Biosciences |
| | Arg1 | A1exF5 | ThermoFisher |
| | iNos | CXNFT | ThermoFisher |
